## Supplementary for "Natural gradient Bayesian sampling automatically emerges in canonical cortical circuits": 1d.py

```
#!/usr/bin/env python
from cann_simulator import *
from analysis_tools import *
from plot_data import *
import warnings
warnings.filterwarnings('ignore')
if __name__ == "__main__":
    params = {
                'time_constant_exc': 1.0,
                'position_max': 180.0,
                'position_min': -180.0,
                'gaussian_width_exc': 40.0,
                'gaussian_width_ES': 20.0,
                'num_neurons': 180,
                'simulation_time': 500.0,
                'time_step': 0.01,
                'recording_start': 50,
                'Fano_factor': 0.5,
                'normalization_k': 0.0005,
                'inhibitory_gain': 10,
                'input_position': 0,
                'feedforward_scale': 0.71648,
                't_steady':50,
                'initial_mean_eq':0,
                'initial_var_eq':30,
                'initial_scale_eq':1e-1
            }
    # Strongly suggest to run each part seperately, otherwise the simulation time will be too long, you can comment the part you don't want to run
    # You can run Figure 1 in one go
    # I highly recommend to run the part of Figure 2 seperatly, in each subfigure, I will provide the parameters you can adjust in a block
    
    #Figure D E possion ff  + likelihood
    plot_possion_ff_likelihood(params,I =6)
    # Show the simulation result for one run CANN model
    # Figure 1 F G
    # plot_possion_ff_likelihood(params,I =6)
    plot_one_simulation_ham(params = params, Rf = 10 ,test_eq = 'normal',tstart=50,tend=60)
    # Figure 1 H
    plot_bump_position_EI(params, Rf = 10 ,test_eq = 'normal',sampling='Hamiltonian')
    
    # Langevin simulation
    # Figure 2 B precision vs Rf 4
    Rf_list = np.linspace(1,16,15)
    plot_precision_vs_Rf(params,ff_scale= params['feedforward_scale'],Rf_list=Rf_list,sampling='Langevin',n_trials=1)
    
    # Figure 2 C bump_height_vs_fisher_information
    Wee_list = np.linspace(0,0.5,2)
    Rflist1 = np.linspace(0.5, 4, 14)
    Rflist2 = np.linspace(4, 6, 4)

    Rf_list = np.concatenate((Rflist1, Rflist2))
    plot_bump_height_vs_Rf(params, Wee_list, Rf_list, test_eq="normal")  

    # Figure 2 D bump_height_vs_time_const
    list1 = np.linspace(0, 0.5, 2)
    list2 = np.linspace(0.51, 0.52, 5)
    list3 = np.linspace(0.53, 0.61, 4)
    list3 = np.linspace(0.62, 1, 11)
    Wee_list = np.concatenate((list1, list2, list3))
    plot_bump_height_vs_time_const(params, Wee_list, Rf = 2, num_trials=100,test_eq="normal")
    
    # Strongly suggest to run them seperately
    # Figure 2 EGH the kl div and cross correlation
    num_trials = 500
    # Figure 2 E plot kl div with different Wee at equilibrium
    Get_Kl_div_vs_Wee(params = params, Wee_list = np.linspace(0, 1, 3),Rf = 3, num_trials=num_trials ,test_eq = 'eq')
    
    # Figure 2 GH plot kl div as input intensity Rf change when Wee = 0 at equilibrium and also the cross correlation
    # show the decaying speed is same as natural gradient Langvelin sampling
    Get_Kl_div_vs_Rf(params = params, Wee = 0, num_trials=num_trials ,test_eq = 'eq')
    
    # Figure 2 F plot kl div convegence time vs RF
    # This one takes a day on 512GB biohpc, but it will save the data
    plot_conv_vs_Rf(params, Wee_list, Rf_list,num_trials=500, test_eq="eq",data_dir="nt500")
    # If you have already run the above code, you can load the data and it will plot it and also run the missing data
    # plot_conv_vs_Rffromdic(params, Wee_list, Rf_list,num_trials=500, test_eq="eq",data_dir="nt500",load_saved=True)
    
    # Figure 2 IJK non equilibrium anneeling strategy
    params['simulation_time'] = 100
    params['recording_start'] = 20
    params['t_steady'] = 20
    plot_one_simulation_Lan(params = params, Rf = 3 ,test_eq = 'non-eq')
    # Figure 2 K plot kl div with different Wee out of equilibrium
    Get_Kl_div_vs_Wee(params = params, Rf = 3, num_trials= 100,test_eq = 'non-eq')

    # Figure 4 c precision vs Rf
    simulator = CANNSimulator(params)#,seed=42)
    simulator.initialize_network()
    Rf_list = np.linspace(1,16,15)
    Wc = simulator.params['critical_weight']
    plot_precision_vs_Rf(params,ff_scale=1.8*Wc ,Rf_list=Rf_list,sampling='Hamiltonian',n_trials=10)
    

    # Figure S1
    plot_one_attractor(params, Rf=10,test_eq="normal",tstart = 0,tend = 100)
    plot_tuning_curves(n_neurons=9, variance=40, sigma=40)
    #Figure S3 wef wei manifold
    simulator = CANNSimulator(params)
    simulator.initialize_network()
    Wc = simulator.params['critical_weight']
    rho = simulator.params['neuron_density']
    ffm = 0.1
    ffx= 4
    size = 101
    fn = f'jsresults_ff_from{ffm}_to{ffx}_with{size}kld.csv'
    wei_list = np.linspace(0, 2, size) * Wc
    ff_list = np.linspace(ffm, ffx, size) * Wc 
    get_js_div_ham(params, wei_list=wei_list,ff_list=ff_list,output_file=fn)
    marker = (1.8*Wc, 0.6*Wc,"red")
    plot_jsd_heatmap_with_min_line(csv_path=fn,rho =rho, critical_weight=Wc,markers=[marker])
```
