## Supplementary for "Natural gradient Bayesian sampling automatically emerges in canonical cortical circuits": analysis_tools.py

```
import numpy as np
from tqdm import tqdm
from sklearn.linear_model import LinearRegression
from cann_simulator import CANNSimulator
import multiprocessing as mp
import pandas as pd
from functools import partial
from scipy.optimize import curve_fit
import gc
import os
def compute_kl_divergence(param_dict,Se,Rf,num_trials = 50):
    """
    Computes the Kullback-Leibler divergence betWeen sampling from CANN and reference distributions from feedforward input.    
    Returns:
        kl: KLD values over time.
    """

    # Load parameters
    rho = param_dict['num_neurons'] / ( param_dict['position_max'] - param_dict['position_min'])
    a = param_dict['gaussian_width_exc']
    vth = 1/(np.sqrt(2 * np.pi) * rho * Rf / a)
    mth =  param_dict['input_position']
    # Compute KL divergence over time
    kl = []
    T = param_dict["simulation_time"]
    dt = param_dict["time_step"]
    counter = np.arange(1, Se.shape[1]+1, 1)
    kl_all = []
    for i in tqdm(counter, desc="Computing KL divergence"):
        Sshort = Se[:, :i].reshape(-1)
        SeMean = np.mean(Sshort)
        SeVar = np.var(Sshort)
        SeVar = max(SeVar, 1e-10)  # Avoid division by zero
        kl = 0.5 * (np.log(vth / SeVar) + (SeVar + (SeMean - mth)**2) / vth - 1)
        kl_all.append(kl)

    return np.array(kl_all)
def compute_cross_correlation(se,timesteps = 2000):
    """
    Computes the cross-correlation for each method in the simulation data.

    Args:
        se_dict: Dictionary containing simulation data for different methods.

    Returns:
        A dictionary with methods as keys and cross-correlations as values.
    """

    correlations = []
    for sim in se:
        sim_trimmed = sim[timesteps:]
        cc = np.correlate(sim_trimmed, sim_trimmed, mode='full')
        cc = cc[cc.size // 2:]  # Keep non-negative lags
        cc = cc / np.max(cc)    # Normalize
        correlations.append(cc)

    # Pad to the same length if needed
    max_len = max(len(c) for c in correlations)
    padded = [np.pad(c, (0, max_len - len(c)), 'constant') for c in correlations]

    cross_correlations = np.mean(padded, axis=0)
    return cross_correlations


def exponential_decaycc(tau, A, tau_c):
    return A * np.exp(-tau / tau_c)

def extract_time_constants(cc):
    """
    Fit exponential decay to cross-correlations to extract time constants.
    """
    tau = np.arange(len(cc))

    # Use only part of the curve (e.g., where decay is smooth)
    cutoff = np.argmax(cc < 0.05) if np.any(cc < 0.05) else len(cc)
    fit_tau = tau[:cutoff]
    fit_cc = cc[:cutoff]

    try:
        popt, _ = curve_fit(exponential_decaycc, fit_tau, fit_cc, p0=(1.0, 100.0))
        _, tau_c = popt
        time_constants = tau_c
    except RuntimeError:
        time_constants = np.nan  # Fit failed
    return time_constants


def exponential_decaykl(t, a, tau, c):
    """
    a * exp(-t/tau) + c
    """
    return a * np.exp(-t / tau) + c


def get_convergence_time(kl, start_index=2, threshold_ratio=0.05, trim_tail=True):
    """
    Computes the convergence time for a KL-divergence sequence, optionally trimming
    small noisy tail segments before fitting or threshold detection.

    Args:
        kl (array-like): sequence of KL values over time
        start_index (int): number of initial points to skip (transient)
        threshold_ratio (float): fraction of the peak value defining "converged"
        trim_tail (bool): whether to remove the trailing segment below threshold before fitting

    Returns:
        int: index (in the original series) at which KL first falls below threshold,
             or np.nan if the series is too short.
    """
    kl = np.asarray(kl)
    n = kl.size
    if n <= start_index:
        return np.nan

    # Post-transient segment
    seg = kl[start_index:]
    peak = seg.max() if seg.max() != 0 else 1.0
    thresh = threshold_ratio * peak

    # Identify tail start: first point below threshold
    below = np.where(seg < thresh)[0]
    if trim_tail and below.size > 0:
        # Trim segment up to first crossing, discarding deeper noisy tail
        fit_seg = seg[:below[0]]
        t_fit = np.arange(fit_seg.size)
    else:
        fit_seg = seg
        t_fit = np.arange(seg.size)

    # Fit exponential decay to trimmed segment to extract time constant
    try:
        p0 = (peak, 1.0, seg.min())  # (amplitude, tau, offset)
        popt, _ = curve_fit(exponential_decaykl, t_fit, fit_seg, p0=p0)
        tau = popt[1]
    except (RuntimeError, ValueError):
        tau = np.nan

    # Find convergence index in full segment
    if below.size > 0:
        conv_idx = below[0] + start_index
    else:
        conv_idx = n - 1

    return conv_idx

def simulate_bump_dynamics(simulator, Rf, Wee, num_trials, test_eq):
    """
    Run the CANN simulator and return the steady‐state bump positions.
    """
    results = simulator.compute_bump_positions_height_over_trials(
        Rf,
        Wei=0,
        ff_scale=simulator.params['feedforward_scale'],
        Wee=Wee,
        num_trials=num_trials,
        test_eq=test_eq
    )
    t0 = int(simulator.params['t_steady'] / simulator.params['time_step']) - 1
    return results['bump_positions'][:, t0:]

def compute_conv_times_for_Wee(params, simulator, Wee, Rf_list, num_trials, test_eq, save_dir=None):
    """
    For a fixed Wee, compute convergence times over all Rf in Rf_list.
    If Wee == 0, also compute the 'fisher' variant.
    Optionally save 'bumps' to file for each Rf.
    """
    conv = []
    conv_fisher = []

    if save_dir:
        os.makedirs(save_dir, exist_ok=True)

    for Rf in tqdm(Rf_list, desc=f"Wee={Wee:.3f}"):
        # 1) simulate
        bumps = simulate_bump_dynamics(simulator, Rf, Wee, num_trials, test_eq)

        # Optionally save bumps
        if save_dir:
            file_name = f"bumps_Wee{Wee:.3f}_Rf{Rf:.3f}_nt{num_trials}.npy"
            np.save(os.path.join(save_dir, file_name), bumps)

        # 2) compute KL & convergence
        kl = compute_kl_divergence(params, bumps, Rf, num_trials=num_trials)
        conv.append(get_convergence_time(kl))

        # 3) extra fisher‐style run only when Wee == 0
        if Wee == 0:
            initials = bumps[:, 0]
            samp = langevin_sampling(
                params, Rf, initials, num_trials=num_trials,
                test_eq=test_eq, conditioner=True
            )
            klf = compute_kl_divergence(params, samp, Rf, num_trials=num_trials)
            conv_fisher.append(get_convergence_time(klf))
            fisher_file = os.path.join(save_dir, f"samp_fisher_Rf{Rf:.3f}_nt{num_trials}.npy")
            np.save(fisher_file, samp)
            del samp, initials
            gc.collect()

    return np.array(conv), np.array(conv_fisher)


def Lan_run_trial(T, dt, rate, precision, mean, initial_state):
    """Function must be at the top level for pickling."""
    num_steps = int(T/dt)+1
    trial = np.zeros(num_steps)
    trial[0] = initial_state

    #trial[0] = np.random.normal(0,1)
    # noise = np.random.normal(0, 1, num_steps-1)

    for t in range(1, num_steps):
        trial[t] = trial[t-1] + rate * precision * (-trial[t-1] + mean) * dt + np.sqrt(2 * rate * dt) * np.random.normal(0,1)#noise[t-1]
        # trial[t] = trial[t-1] + rate * precision * (-trial[t-1] + mean) * dt + np.sqrt(2 * rate * dt) * noise[t]
    return trial

def langevin_sampling(param_dict, Rf,initial_states_list, num_trials=50, test_eq=False, conditioner=True):
    # np.random.seed(seed)  # Set seed for reproducibility

    # Load parameters
    # if test_eq:
        
    # else:
    #     T = param_dict["simulation_time"]
    T = param_dict["simulation_time"]-param_dict['t_steady']
    tau = param_dict["time_constant_exc"]
    dt = param_dict["time_step"]
    ff_scale = param_dict["feedforward_scale"]
    
    rho = param_dict['num_neurons'] / (param_dict['position_max'] - param_dict['position_min'])
    a = param_dict['gaussian_width_exc']
    precision = (np.sqrt(2 * np.pi) * rho * Rf) / a
    mean = param_dict['input_position']

    # OU process parameters
    if conditioner:
        rate =  a * ff_scale/ (rho* tau * Rf * np.sqrt(2 * np.pi))
    else:
        rate = a * ff_scale / (rho* tau * np.sqrt(2 * np.pi))
    # Use `partial` to pre-fill arguments
    run_trial_partial = partial(Lan_run_trial, T, dt, rate, precision, mean)

    # Run trials in parallel
    with mp.get_context('spawn').Pool() as pool:
        desc = "Natural Gradient Langevin sampling" if conditioner else "Pure Langevin sampling"
        results = list(tqdm(
            pool.map(run_trial_partial, initial_states_list),  # Pass only `i` (trial index)
            total=num_trials,
            desc=desc
        ))

    return np.array(results)


def compute_jsd(Iprec, SeVar, SeMean):
    mth = 0
    vth = 1 / Iprec

    mm = 0.5 * (mth + SeMean)
    mv = 0.5 * (vth + SeVar) - (mth * SeMean)

    # KL divergence between (mv, mm) and (SeVar, SeMean)
    KL1 = 0.5 * (np.log(mv / SeVar) + (SeVar + (SeMean - mm)**2) / mv - 1)

    # KL divergence between (vth, mth) and (SeVar, SeMean)
    
    KL2 = 0.5 * (np.log(vth / SeVar) + (SeVar + (SeMean - mth)**2) / vth - 1)

    # JSD = KL2
    JSD = 0.5 * KL1 + 0.5 * KL2

    return JSD

def process_one_combination(args):
    simulator_params, wei, ff, Rf, Iprec = args
    simulator = CANNSimulator(simulator_params)
    simulator.initialize_network()
    tr = simulator.params['recording_start']
    dt  = simulator.params['time_step']
    rho = simulator.params['neuron_density']
    re = simulator.get_bump_position(
        Rf=Rf,
        Wei=-wei,
        ff_scale= ff,
        Wee=simulator.params['recurrent_weight_e2e'],
        test_eq='normal'
    )
    re = re[int(tr/dt):]
    SeMean = np.mean(re)
    SeVar = np.var(re)
    jsd = compute_jsd(Iprec, SeVar, SeMean)
    
    return {'Wei': wei, 'ff': ff, 'SeMean': SeMean, 'SeVar': SeVar, 'JSD': jsd}

def get_js_div_ham(params,wei_list = np.linspace(0, 2, 10),
    ff_list = np.linspace(0.2, 3, 10) ,output_file='results.csv'):
    simulator = CANNSimulator(params)
    simulator.initialize_network()
    
    rho = simulator.params['neuron_density']
    a = simulator.params["gaussian_width_exc"]
    critical_weight = simulator.params['critical_weight']
    

    # ff_list = rho / np.sqrt(2) * ff_list
    
    Rf = 10
    Iprec = (np.sqrt(2 * np.pi) * rho / a * Rf)
    
    task_args = [(params, wei, ff, Rf, Iprec) for wei in wei_list for ff in ff_list]

    with mp.Pool() as pool:
        results = pool.map(process_one_combination, task_args)
    
    df = pd.DataFrame(results)
    df.to_csv(output_file, index=False)
```
