## Supplementary for "Natural gradient Bayesian sampling automatically emerges in canonical cortical circuits": cann_simulator.py

```
import numpy as np
import numpy.matlib
import matplotlib.pyplot as plt
from scipy.stats import multivariate_normal
from matplotlib.collections import LineCollection
from tqdm import tqdm
import pickle
import multiprocessing as mp
from functools import partial
import sys
from collections import defaultdict
class CANNSimulator:
    def __init__(self, params=None):
        """
        Initialize the CANNSimulator with given parameters.

        Parameters:
        params (dict, optional): Dictionary of parameters to override default values.
        
    A simulator for Continuous Attractor Neural Network (CANN) models.

    This class provides methods to initialize network parameters, run simulations,
    and analyze the results of CANN models. It supports various configurations
    and allows for the calibration of feedforward weights.

    Attributes:
        params (dict): Dictionary of simulation parameters.
        conn_params (dict): Dictionary of connection parameters.
        rng (np.random.Generator): Random number generator for reproducibility.

    Methods:
        initialize_network(): Initialize network connectivity.
        run_simulation(Rf, Wei, ff_scale, Wee=None, seed=None): Run the main simulation.
        calibrate_ff_scale(Rf_range=(0.1, 2), num_points=20): Calibrate feedforward weight.
        plot_precision_vs_Rf(Rf_range=(0.1, 2), num_points=2): Plot precision vs input intensity.
        analyze_results(results, Rf): Analyze simulation results.
        plot_results(results, analysis): Plot simulation results.
        compute_bump_positions_height_over_trials(Rf, Wei, ff_scale, num_trials=50, Wee=None): Compute bump positions and height over multiple trials.
        run_sim_vs_Wee_Rf(Wei, ff_scale, num_trials=50, Wee_list=None, Rf_list=None): Run simulations for different Wee and Rf values.
    """
    def __init__(self, params=None):

        default_params = {
            'time_constant_exc': 1.0,
            'position_max': 180.0,
            'position_min': -180.0,
            'gaussian_width_exc': 40.0,
            'gaussian_width_ES': 20.0,
            'num_neurons': 180,
            'simulation_time': 100.0,
            'time_step': 0.01,
            'recording_start': 20,
            'Fano_factor': 0.5,
            'normalization_k': 0.0005,
            'inhibitory_gain': 10,
            'input_position': 0,
            'feedforward_scale': 0.58214787,
            't_steady':20,
            'initial_mean_eq':0,
            'initial_var_eq':60,
            'initial_scale_eq':1e-1
        }
        self.params = {**default_params, **(params or {})}
        self._compute_parameters()
        self.conn_params = None
        # self.rng = np.random.default_rng(seed)
        
    def _compute_parameters(self):
        """Initialize and compute all simulation parameters"""
        
        p = self.params
        #
        p['time_constant_som'] = p['time_constant_exc']

        p['gaussian_width_SE'] = np.sqrt(p['gaussian_width_exc'] ** 2 - p['gaussian_width_ES'] ** 2)

        p['gaussian_width_SS'] = np.sqrt( (p['gaussian_width_exc'] ** 2 + p['gaussian_width_ES'] ** 2) / 2)

         # Position range = max - min
        p['position_range'] = p['position_max'] - p['position_min']
        
        # Neuron density = number / range
        p['neuron_density'] = p['num_neurons'] / p['position_range']
        
        #
          # Generate preferred stimulus points for neurons
        p['PrefStim'] = np.linspace(
            p['position_min'], p['position_max'], p['num_neurons'] + 1
        )[1:p['num_neurons'] + 1]
        
        # Maximum inhibition strength
        p['max_inhibition'] = 1 / (12 * p['neuron_density'])
        
        # Critical weight calculation
        numerator = 8 * np.sqrt(2 * np.pi) * p['normalization_k'] * p['gaussian_width_exc']
        p['critical_weight'] = np.sqrt(numerator / p['neuron_density'])

      
        # Recurrent connection weights
        p['recurrent_weight_e2e'] = 0.5 * p['critical_weight']  # E->E
        p['recurrent_weight_e2i'] = 0.5 * p['critical_weight']  # E->I
        
        # Feedforward intensity calculation
        denominator = 2 * np.sqrt(np.pi) * p['normalization_k'] * p['gaussian_width_exc']
        p['feedforward_intensity_scale'] =  (p['critical_weight'] / denominator) #Ufd
        

    def initialize_network(self):
        """Initialize network connectivity"""

        param_dict = self.params
        PrefStim = param_dict["PrefStim"]
        a_SE = param_dict["gaussian_width_SE"]
        xrange = param_dict['position_range']

        aE = param_dict["gaussian_width_exc"]

        a_ES = param_dict['gaussian_width_ES']


        ## Connection kernels
        W_angle = np.angle(np.exp((1j*(PrefStim-PrefStim[0])*(2*np.pi/xrange)))) * xrange/(2*np.pi)
        W_kerFtE = np.exp(-W_angle**2/(2*aE**2))/(np.sqrt(2*np.pi)*aE)    
        WE = np.expand_dims(W_kerFtE,axis=1)
        WE = np.fft.fft(W_kerFtE)
        
        W_kerFtES = np.exp(-W_angle**2/(2*a_ES**2))/(np.sqrt(2*np.pi)*a_ES)   
        WES = np.expand_dims(W_kerFtES,axis=1) 
        WES = np.fft.fft(W_kerFtES)
        
        W_kerFtSE = np.exp(-W_angle**2/(2*a_SE**2))/(np.sqrt(2*np.pi)*a_SE) 
        WSE = np.expand_dims(W_kerFtSE,axis=1)   
        WSE = np.fft.fft(W_kerFtSE)
        
        # figc1, axc1 = plt.subplots(figsize=(8,8))
        # figc1.subplots_adjust(left=0.15, bottom=0.2, right=0.85, top=0.9, wspace=0.04, hspace=0.3)
        # axc1.plot(PrefStim,W_kerFtE, label="Recurrent Kernel",linewidth=1)
        # axc1.plot(PrefStim,W_kerFtES, label="WES Kernel",linewidth=1)
        # axc1.plot(PrefStim,W_kerFtSE, label="WSE Kernel",linewidth=1)
        # axc1.legend()

        
        self.conn_params =  {"E Kernel": WE,'ES Kernel':WES, 'SE Kernel':WSE}


    def _temporal_ff(self, Rf,T,mu):
        """Calculate temporal feedforward input"""
        a = self.params["gaussian_width_exc"]
        xrange = self.params['position_range']
        PrefStim = self.params['PrefStim']
        # mu = self.params['input_position']
        # T = self.params['simulation_time']
        dt = self.params['time_step']
        N = self.params['num_neurons']
        x1 =  np.repeat(mu,N)    
        pos1 = np.subtract(x1,PrefStim)
        pos1 = np.angle(np.exp(1j*pos1 * (2*np.pi)/xrange)) * xrange/(2*np.pi)
        pos1 = np.expand_dims(pos1,axis=1)
        pos1 = np.matlib.repmat(pos1,1,int(T/dt))
        Ipos1 = Rf * np.exp(-(pos1**2) / (4*a**2))
        #print(np.shape(Ipos1))

        return Ipos1
    
    def run_simulation(self, Rf, Wei, ff_scale, Wee,test_eq="normal",noise = True):
        """Main simulation function ff_scale = Wef *rho / (sqrt(2))"""
        param_dict = self.params
        N = param_dict["num_neurons"]
        T = param_dict["simulation_time"]
        dt = param_dict["time_step"]
        # tau = param_dict["time_constant"]
        a = param_dict["gaussian_width_exc"]
        aS = param_dict["gaussian_width_SS"]
        k = param_dict["normalization_k"]
        gain = param_dict["inhibitory_gain"]
        rho = param_dict["neuron_density"]
        F = param_dict["Fano_factor"]
        mu = param_dict['input_position']
        t_steady = param_dict['t_steady']
        tauE = param_dict["time_constant_exc"]
        tauS = param_dict['time_constant_som']
        WE = self.conn_params['E Kernel']
        WES = self.conn_params['ES Kernel']
        WSE = self.conn_params['SE Kernel']
        Wie = param_dict["recurrent_weight_e2i"]

        # print("1",test_eq)
        # Initialize arrays
        U = np.zeros([N,int(T/dt)])
        V = np.zeros([N,int(T/dt)])
        Oc = np.zeros([N, int(T/dt)])
        Vc = np.zeros([N, int(T/dt)])

        # U[:,0] =  np.random.normal(0,1,N)
        # V[:,0] =  np.random.normal(0,1,N)
        # Oc[:,0] = np.random.normal(0,1,N)
        # Vc[:,0] =  np.random.normal(0,1,N)
        # if not noise:
        #     F = 0
        if test_eq == 'eq':        
            initial_mean = param_dict['initial_mean_eq']
            initial_var = param_dict['initial_var_eq']
            
            shift = np.random.normal(initial_mean,initial_var) #60
            Iext1 = self._temporal_ff(Rf,t_steady,int(shift)/2*2)
            Iext2 = self._temporal_ff(Rf,T-t_steady,mu)
            Iext = np.concatenate((Iext1,Iext2),axis=1)
            # print(Iext[:,1])
            # print(Iext[1,:])
        elif test_eq == 'non-eq':
            initial_scale = param_dict['initial_scale_eq']

            small_input =initial_scale  #Rf*initial_scale #
            Iext1 = small_input*np.ones([N,int(t_steady/dt)])
            Iext2 = self._temporal_ff(Rf,T-t_steady,mu)
            Iext = np.concatenate((Iext1,Iext2),axis=1)
        else:
            Iext = self._temporal_ff(Rf,T,mu)

        # if Wee is None:
        #     Wee = param_dict["recurrent_weight_e2e"]
        # if seed is not None:
        #     np.random.seed(seed)
        #np.random.seed(1)
        # Run simulation
        for i in range(int(T/dt) - 1):
            # Noise terms (clip negatives to zero)
            noisevr = U[:,i].clip(min=0)
            noiseI = V[:,i].clip(min=0)
            
            # Firing Rates for this time point
            O = Oc[:,i]  # No np.array() needed
            Oin = Vc[:,i]
            
            FRE = np.real(np.fft.ifft(WE * np.fft.fft(O)))
            #FRE = np.roll(FRE, int(len(FRE)/2))
            FRSE = np.real(np.fft.ifft(WSE * np.fft.fft(O)))
          #  FRSE = np.roll(FRSE, int(len(FRSE)/2))
            FRI = np.real(np.fft.ifft(WES * np.fft.fft(Oin)))
           # FRI = np.roll(FRI, int(len(FRI)/2))
            
            U[:,i+1] = ((1-dt/tauE)* U[:,i]) + (Wee*FRE*dt/tauE) + (Wei*FRI*dt/tauE) + (ff_scale*Iext[:,i]*dt/tauE) + (np.sqrt(F*noisevr*dt/tauE) * np.random.normal(0,1,N))
            V[:,i+1] = ((1-dt/tauS)* V[:,i]) + (Wie*FRSE*(dt/tauS)) #+ (np.sqrt(F*noiseI*dt) * np.random.normal(0,1,N))

            # Divisive normalization
            Ocn = U[:,i+1].clip(min=0) ** 2
            Ocd = np.multiply(k*rho, np.sum(Ocn, axis=0))
            Oc[:,i+1] = Ocn / (Ocd + 1)
            
            # Inhibitory
            Vc[:,i+1] = gain * V[:,i+1]
        
        # Optional: Final check (likely redundant)
        Oc = Oc.clip(min=0)
        
        return {
            'E_stim': self._calculate_stimulus(Oc),
            'I_stim': self._calculate_stimulus(Vc),
            'E_bump_height': np.sum(U, axis=0) / (np.sqrt(np.pi) * 2* a * rho),
            'I_bump_height': np.sum(V, axis=0) / (np.sqrt(np.pi) * 2* aS * rho),
            'E_FR': Oc,
            'I_FR': Vc,
            'Synaptic_Input': U,
            'I_Synaptic_Input': V,
            'params': self.params
        }
         
    def _calculate_stimulus(self, activity):
        """Calculate stimulus position from activity"""
        pref_factor = self.params['PrefStim'] * np.pi / 180
        if np.all(activity == 0):
            return np.zeros(activity.shape[1])
        e = np.exp(1j * pref_factor) @ activity
        return np.angle(e, deg=True)
    
    def calibrate_ff_scale(self, Rf_list=np.linspace(1,25,23),  Wei=0,n_trials = 5000):
        """Calibrate feedforward weight using precision analysis"""
        Ufd = self.params['feedforward_intensity_scale']
        #Rf_list = np.linspace(*Rf_range, num_points)*Ufd
        t_record = self.params['recording_start']
        dt = self.params['time_step']
        rho = self.params['neuron_density']
        a = self.params["gaussian_width_exc"]
        
        record_precision_inv = []
        
        # Create partial function with fixed arguments
        #simulate_trial = partial(self.run_simulation, Wei=Wei, ff_scale=1, Wee=self.params['recurrent_weight_e2e'], test_eq = 'normal')
        
        simulate_trial = partial(
            self.get_bump_position,
            Wei=Wei,
            ff_scale=self.params['feedforward_scale'],
            Wee=self.params['recurrent_weight_e2e'],
            test_eq='normal'
        )

        # Define number of trials per Rf
        

        # Expand the Rf_list: for each Rf, repeat it n_trials times
        expanded_Rf_list = [Rf for Rf in Rf_list for _ in range(n_trials)]

        # Optional: you might want to include trial info, e.g., (Rf, trial_num) for more control
        # expanded_Rf_list = [(Rf, i) for Rf in Rf_list for i in range(n_trials)]
        # simulate_trial would then need to accept (Rf, trial_num) and handle accordingly

        # Run simulations in parallel
        with mp.get_context('spawn').Pool() as pool:
            results = list(tqdm(
                pool.imap(simulate_trial, expanded_Rf_list),
                total=len(expanded_Rf_list),
                desc=f"Running {n_trials} trials per Rf", file=sys.stdout)
            )

        # --- after running `results = pool.imap(...)` with expanded_Rf_list ---

        # 1) Compute 1/variance per trial
        record_precision_inv = []
        for res in results:
            Se = res.flatten()
            Se = Se[int(t_record/dt):]
            record_precision_inv.append(1.0 / np.var(Se))

        # 2) Group into blocks of size n_trials and average
        mean_precision_inv = []
        for i, Rf in enumerate(Rf_list):
            block = record_precision_inv[i * n_trials : (i + 1) * n_trials]
            mean_precision_inv.append(np.mean(block))

        # 3) Linear regression on mean precisions
        slope = np.polyfit(Rf_list, mean_precision_inv, 1)[0]

        # 4) Compute theoretical slope and update ff_scale
        theoretical_slope = np.sqrt(2 * np.pi) * rho / a
        new_ff = theoretical_slope / slope
        self.params['feedforward_scale'] = new_ff

        # 5) Rerun with updated ff_scale (same expanded_Rf_list)
        simulate_trial = partial(
            self.get_bump_position,
            Wei=Wei,
            ff_scale=new_ff,
            Wee=self.params['recurrent_weight_e2e'],
            test_eq='normal'
        )
        with mp.get_context('spawn').Pool() as pool:
            new_results = list(tqdm(
                pool.imap(simulate_trial, expanded_Rf_list),
                total=len(expanded_Rf_list),
                desc="Recalculating precision with updated ff_scale", file=sys.stdout
            ))

        # 6) Compute new mean precision per Rf
        record_precision_inv_new = []
        for res in new_results:
            Se = res.flatten()
            Se = Se[int(t_record/dt):]
            record_precision_inv_new.append(1.0 / np.var(Se))

        mean_precision_inv_new = []
        for i, Rf in enumerate(Rf_list):
            block = record_precision_inv_new[i * n_trials : (i + 1) * n_trials]
            mean_precision_inv_new.append(np.mean(block))

        # 7) Plot
        plt.figure(figsize=(10, 6))
        plt.plot(Rf_list,
                theoretical_slope * np.array(Rf_list),
                label='Theoretical Prediction')
        plt.plot(Rf_list,
                mean_precision_inv_new,
                'o',
                label='Simulation (mean over 80 trials)')
        plt.xlabel('Feedforward Intensity (Rf)')
        plt.ylabel('1 / Variance (Precision)')
        plt.title('Precision vs Input Intensity')
        plt.legend()
        plt.grid(False)
        plt.savefig(f'precision_vs_Rf_wei{Wei}_ff{new_ff:.3f}.svg')

        return new_ff
    def get_bump_position(self, Rf, Wei, ff_scale, Wee,test_eq):
        result = self.run_simulation(Rf, Wei, ff_scale, Wee,test_eq)
        return result['E_stim']
    
    def _get_bump_height_position(self, Rf, Wei, ff_scale, Wee,test_eq): 
        result = self.run_simulation(Rf, Wei, ff_scale, Wee,test_eq)
        return {'E_bump_height':result['E_bump_height'], 'E_stim':result['E_stim']}

    def compute_bump_positions_height_over_trials(self, Rf, Wei, ff_scale,Wee, num_trials=50,test_eq = 'eq'):
        """Computes excitatory stimulation for input conditions over multiple trials.
    The default input conditions are for langevin sampling.
  """

        args = [
            (Rf, Wei, ff_scale, Wee, test_eq)
            for i in range(num_trials)
        ]
        # print(test_eq)
        with mp.get_context('spawn').Pool() as pool:
            results = list(
                tqdm(
                    pool.starmap(self._get_bump_height_position, args),
                    total=num_trials,
                    desc=f"Running Rf={Rf:.2f}"
                )
            )
        all_trials_pos = [result['E_stim'] for result in results]
        all_trials_height = [result['E_bump_height'] for result in results]

        return {
            'time': self.params['simulation_time'],
            'bump_positions': np.array(all_trials_pos),
            'bump_height': np.array(all_trials_height)
        }


def save_simulation_data_pickle(data, filename):
    with open(filename, 'wb') as f:
        pickle.dump(data, f)   
def load_simulation_data_pickle(filename):
    with open(filename, 'rb') as f:
        return pickle.load(f)

def bump_position_ham(params, Rf=10):
    simulator = CANNSimulator(params)
    simulator.initialize_network()
    results = simulator.run_simulation(
        Rf = Rf,#0.8 * simulator.params['feedforward_intensity_scale'], 
        Wei= -0.6*simulator.params['critical_weight'], 
        ff_scale=1.624,#1.3*simulator.params['critical_weight'],
        Wee = simulator.params['recurrent_weight_e2e']
    )
    ZE = results['E_stim']
    ZS = results['I_stim']
    UE = results['E_bump_height']
    UI = results['I_bump_height']
    return ZE, ZS, UE, UI
```
