## Supplementary for "Natural gradient Bayesian sampling automatically emerges in canonical cortical circuits": plot_data.py

```
import matplotlib.pyplot as plt
import seaborn as sns
import numpy as np
from scipy.stats import norm
from matplotlib.collections import LineCollection
from matplotlib.gridspec import GridSpec
from matplotlib.cm import get_cmap
from cann_simulator import *
from analysis_tools import *
import pandas as pd
from mpl_toolkits.axes_grid1 import make_axes_locatable
import matplotlib.colors as mcolors
import seaborn as sns
import os
from datetime import datetime
import json
import shutil
import itertools
from joblib import Parallel, delayed
from multiprocessing import cpu_count
import warnings
import pickle # For saving/loading scan data


warnings.filterwarnings('ignore')
np.seterr(divide='ignore', invalid='ignore')

import os
import sys
import gc


def save_fig(fig, filename, sampling):
    """
    Save the figure without background.
    Args:
    - fig: The figure to save.
    - filename: The name of the file to save the figure as.
    - sampling: The type of sampling  we refered to in the simulation.
    """
    fig.patch.set_facecolor('none')   # Set the patch (outer part) to transparent
    fig.set_facecolor('none')         # Set the figure facecolor to transparent
    # plt.rcParams['text.usetex'] = True
    # Turn off axis for all subplots
    for ax in fig.axes:
        #ax.set_axis_off()
        #ax.set_xlim(left=0)
        ax.tick_params(
        axis='both',        # Apply to both x and y
        which='both',       # Apply to both major and minor ticks
        direction='in',     # Tick direction: inward
        length=4,
        width=1,
        labelsize=10
        )
        xlabel = ax.get_xlabel()  
        ylabel = ax.get_ylabel()  

        ax.set_xlabel(xlabel, fontsize=16) 
        ax.set_ylabel(ylabel, fontsize=16)

        # Optional: make ticks thinner and closer if desired
        ax.tick_params(width=1, length=4, labelsize=10)

    if sampling == 'Hamiltonian':
        folder_path = 'Ham'
    else:
        folder_path = 'Lan'
    if not os.path.exists(folder_path):
        os.makedirs(folder_path)
    plt.rcParams['ps.fonttype'] = 42
    plt.rcParams['pdf.fonttype'] = 42
    filename = os.path.join(folder_path, filename)
    fig.savefig(filename, format='eps',transparent = True,  bbox_inches='tight')
    plt.close(fig)

def plot_one_simulation_Lan(params, Rf=10,test_eq="normal",tstart = 0,tend = 100):
    """
    Plot one simulation for Langevin sampling

    Args:
    - params: dictionary containing simulation parameters
    - Rf: float, input intensity for Langevin sampling
    - test_eq: str, type of input used for initialization
        -eq: get equilibrium results
        -non-eq: get non-equilibrium results
        -nomal: get normal results/ the initialization is normal
    - tstart: int, start time for plotting
    - tend: int, end time for plotting
    """
    
    simulator = CANNSimulator(params)
    simulator.initialize_network()
    results_Lan = simulator.run_simulation(
        Rf = Rf,#0.8 * simulator.params['feedforward_intensity_scale'], 
        Wei= 0, 
        ff_scale=params['feedforward_scale'],
        Wee = simulator.params['recurrent_weight_e2e'],
        test_eq= test_eq
    )
    plot_results(results_Lan, "Langevin",tstart = tstart,tend = tend)


def plot_one_simulation_ham(params, Rf=10,test_eq="normal",tstart = 0,tend = 100):
    """
    Plot one simulation for Hamiltonian sampling

    Args:
    - params: dictionary containing simulation parameters
    - Rf: float, input intensity for Hamiltonian sampling
    - test_eq: str, type of input used for initialization
        -eq: get equilibrium results
        -non-eq: get non-equilibrium results
        -nomal: get normal results/ the initialization is normal
    - tstart: int, start time for plotting
    - tend: int, end time for plotting
    """
    simulator = CANNSimulator(params)
    simulator.initialize_network()
    results_ham = simulator.run_simulation(
        Rf = Rf,#0.8 * simulator.params['feedforward_intensity_scale'], 
        Wei= -0.6*simulator.params['critical_weight'], 
        ff_scale=1.8*simulator.params['critical_weight'],
        Wee = simulator.params['recurrent_weight_e2e'],
        test_eq= test_eq
    )
    plot_results(results_ham, "Hamiltonian",tstart = tstart,tend = tend)


def plot_results(results,sampling,tstart = 0,tend = 100):
    """Visualization methods"""
    plot_firing_rates(results,sampling,tstart = tstart,tend = tend)    
    plot_bump_position(results, sampling,tstart = tstart,tend = tend)
    plot_bump_height(results,sampling,tstart = tstart,tend = tend)


def plot_firing_rates(results, sampling,tstart = 0,tend = 100):
    """Plot firing rates"""
    fig1,ax1 = plt.subplots(figsize=(2.5,1.75))
    E_FR = results['E_FR'][:,int(tstart/results['params']['time_step']):int(tend/results['params']['time_step'])]
    I_FR = results['I_FR'][:,int(tstart/results['params']['time_step']):int(tend/results['params']['time_step'])]
    plt.imshow(E_FR, aspect='auto', cmap=plt.get_cmap('Blues'),
        extent=[0, tend-tstart, -180, 180], origin='lower', vmin=0, vmax=100)
    ax1.set_yticks([-180, -120,-60,0,60,120,180])
    ax1.set_xlim(left=0)
    plt.colorbar(label='Firing Rate', shrink=0.5, ticks=[0, 50, 100])
    plt.title(f'Excitatory Population Firing Rate for {sampling} Sampling')
    plt.xlabel('Time Steps')
    plt.ylabel('Neuron Index')
    save_fig(fig1, f'one_simulation_FR_all_{sampling}.eps', sampling)
    plt.close()
    if sampling == 'Hamiltonian':
        fig2,ax2 = plt.subplots(figsize=(2.5,1.75))
        
        plt.imshow(I_FR, aspect='auto', cmap=plt.get_cmap('Purples'), extent=[0, tend-tstart, -180, 180], origin='lower', vmin=0, vmax=100)
        ax2.set_yticks([-180, -120,-60,0,60,120,180])
        ax2.set_xlim(left=0)
        plt.colorbar(label='Firing Rate', shrink=0.5, ticks=[0, 50, 100])
        
        plt.title(f'Inhibitory Population Firing Rate for {sampling} Sampling')
        plt.xlabel('Time Steps')
        plt.ylabel('Neuron Index')
        save_fig(fig2, f'one_simulation_FR_inh_all_{sampling}.eps', sampling)
        plt.close()


def plot_bump_position(results, sampling,tstart = 0,tend = 100):
    """Plot bump position over time"""
    t = np.arange(0, tend-tstart, results['params']['time_step'])
    figbp, axbp = plt.subplots(figsize=(2.5,1.75))
    
    axbp.plot(t, results['E_stim'][int(tstart/results['params']['time_step']):int(tend/results['params']['time_step'])],c='blue')
    axbp.set_xlim(0,tend - tstart)
    axbp.set_yticks([-6, -4,-2,0,2,4,6])
    axbp.set_xticks([0,(tend-tstart)/2,tend-tstart])
    axbp.set_xlabel(r'Time /$\tau$')
    axbp.set_ylabel(r'Bump Position')
    axbp.set_title("Bump Position Over Time")
    save_fig(figbp, f'one_simulation_BP_{sampling}.eps', sampling)
    plt.close()
    if sampling == 'Hamiltonian':
        figbp, axbp = plt.subplots(figsize=(2.5,1.75))
        axbp.set_ylim(-6,6)
        axbp.plot(t, results['I_stim'][int(tstart/results['params']['time_step']):int(tend/results['params']['time_step'])],c='purple')
        
        axbp.set_xlim(0,tend - tstart)
        axbp.set_yticks([-6, -4,-2,0,2,4,6])
        axbp.set_xticks([0,(tend-tstart)/2,tend-tstart])
        axbp.set_xlabel(r'Time /$\tau$')
        axbp.set_ylabel(r'Bump Position')
        axbp.set_title("Bump Position Over Time")
        save_fig(figbp, f'one_simulation_BP_Inh_{sampling}.eps', sampling)
        plt.close()

def plot_bump_position_EI(params, test_eq, Rf, sampling):
    """
    Plot the bump position for excitatory-inhibitory network.

    Args:
    - params: dictionary containing simulation parameters
    - Rf: float, input intensity for Hamiltonian sampling
    - test_eq: str, type of input used for initialization
        -eq: get equilibrium results
        -non-eq: get non-equilibrium results
        -nomal: get normal results/ the initialization is normal
    - sampling (str): Sampling identifier.
    """
    
    simulator = CANNSimulator(params)
    simulator.initialize_network()
    results = simulator.run_simulation(
        Rf=Rf,
        Wei=-0.6 * simulator.params['critical_weight'],
        ff_scale=1.8 * simulator.params['critical_weight'],
        Wee=simulator.params['recurrent_weight_e2e'],
        test_eq=test_eq
    )

    # Extract post-recording data
    x1 = results['E_stim'][int(results['params']['recording_start'] / results['params']['time_step']):]
    y1 = results['I_stim'][int(results['params']['recording_start'] / results['params']['time_step']):]

    # Set up 2D Gaussian KDE grid
    x = np.linspace(-6, 6, 200)
    y = np.linspace(-6, 6, 200)
    X, Y = np.meshgrid(x, y)
    pos = np.dstack((X, Y))
    mean = np.mean(np.vstack([x1, y1]).T, axis=0)
    cov = np.cov(np.vstack([x1, y1]))

    rv = multivariate_normal(mean, cov)
    Z = rv.pdf(pos)

    # Create GridSpec layout
    fig = plt.figure(figsize=(6, 6))
    gs = GridSpec(4, 4)

    ax_main = fig.add_subplot(gs[1:, 0:3])
    ax_xDist = fig.add_subplot(gs[0, 0:3], sharex=ax_main)
    ax_yDist = fig.add_subplot(gs[1:, 3], sharey=ax_main)

    # Main heatmap using imshow
    im  = ax_main.imshow(
        Z,
        extent=[x.min(), x.max(), y.min(), y.max()],
        origin='lower',
        cmap='Blues',
        aspect='auto',
        alpha=0.7
    )

    # Trajectory overlay (color by time) 
    trac1 = results['E_stim'][-int((results['params']['time_constant_exc'] * 4) / results['params']['time_step']):]
    trac2 = results['I_stim'][-int((results['params']['time_constant_exc'] * 4) / results['params']['time_step']):]
    ct = np.linspace(0, 1, len(trac1))
    points = np.vstack([trac1, trac2]).T.reshape(-1, 1, 2)
    segments = np.concatenate([points[:-1], points[1:]], axis=1)

    lc = LineCollection(segments, cmap='plasma', norm=plt.Normalize(0, 1), linewidth=2)
    lc.set_array(ct)
    ax_main.add_collection(lc)

    # Marginal histograms
    ax_xDist.hist(x1, bins=40, color='lightsteelblue', alpha=0.6, density=True)
    ax_yDist.hist(y1, bins=40, color='plum', alpha=0.6, density=True, orientation='horizontal')
    ax_xDist.yaxis.set_ticks_position('left')  # Optional: Ensure y-axis is on the left

    ax_xDist.spines['left'].set_visible(True)  # Show left border
    # Marginal KDE plots
    # sns.kdeplot(x=x1, ax=ax_xDist, fill=True, color='steelblue')
    # sns.kdeplot(y=y1, ax=ax_yDist, fill=True, color='purple')
    ax_xDist.plot(x, norm.pdf(x, mean[0], np.sqrt(cov[0, 0])), color='steelblue', linewidth=2)
    ax_yDist.plot(norm.pdf(y, mean[1], np.sqrt(cov[1, 1])), y, color='purple', linewidth=2)

    ax_yDist.xaxis.set_ticks_position('bottom')  # Optional: Ensure x-axis is at the bottom
    ax_yDist.spines['bottom'].set_visible(True)  # Show bottom border
    ax_xDist.set_yticks([0, 0.15, 0.3])
    ax_xDist.set_ylim(0,0.4)
    ax_yDist.set_xticks([0, 0.15, 0.3])
    ax_yDist.set_xlim(0,0.4)
    ax_xDist.set_xlim(ax_main.get_xlim())
    ax_yDist.set_ylim(ax_main.get_ylim())
    plt.setp(ax_xDist.get_xticklabels(), visible=False)
    plt.setp(ax_yDist.get_yticklabels(), visible=False)
    ax_xDist.tick_params(bottom=False)
    ax_yDist.tick_params(left=False)

    # Main plot settings
    ax_main.set_xlim(-6, 6)
    ax_main.set_ylim(-6, 6)
    
    fig.suptitle("Network sampling distribution", fontsize=14)
    ax_main.set_xlabel("Stimulus sample $z_E$ (E neurons)")
    ax_main.set_ylabel("$z_S$ (SOM neurons)")
    cbar_ax = fig.add_axes([0.15, 0.08, 0.7, 0.03])  # [left, bottom, width, height]
    fig.colorbar(im, ax=ax_main, fraction=0.046, pad=0.04)
    cbar = fig.colorbar(lc, cax=cbar_ax, orientation='horizontal')
    cbar.set_label('Elapsed time')
    cbar.set_ticks([0, 1])
    cbar.set_ticklabels(['0', r'10 $\tau$'])

    plt.tight_layout(rect=[0, 0, 1, 0.95])  # Leave some space at the top

    save_fig(fig, f'one_simulation_EI3_BP_{sampling}.eps', sampling)
    plt.close()

def plot_bump_height(results, sampling,tstart = 0,tend = 100):
    """Plot bump height over time"""
    t = np.arange(0, tend-tstart, results['params']['time_step'])
    figbh, axbh = plt.subplots(figsize=(2.5,1.75))
    figbh.subplots_adjust(left=0.1, bottom=0.3, right=0.8, top=0.9, wspace=0.04, hspace=0.3)
    axbh.plot(t, results['E_bump_height'][int(tstart/results['params']['time_step']):int(tend/results['params']['time_step'])], label='Bump Height')
    axbh.set_xlim(0,tend-tstart)
    plt.title(f'Bump Height Over Time for {sampling} Sampling')
    plt.xlabel(r'Time /$\tau$') 
    plt.ylabel(r'Bump Height')
    save_fig(figbh, f'one_simulation_bump_height_{sampling}.eps', sampling)
    plt.close()


def plot_precision_vs_Rf(params,Rf_list=np.linspace(1,25,23),sampling='Langevin',n_trials=80):
    """
    Plot the precision vs feedforward intensity for a given set of parameters.

    Args:
    - params: dictionary containing simulation parameters
    - Rf_list: input intensity for  sampling, array-like, optional, default is np.linspace(1,25,23)
    - sampling: str, optional, default is 'Langevin'
    - n_trials: int, optional, default is 80
    """
    simulator = CANNSimulator(params)
    simulator.initialize_network()

    t_record = simulator.params['recording_start']
    dt       = simulator.params['time_step']
    rho      = simulator.params['neuron_density']
    a        = simulator.params["gaussian_width_exc"]

    theoretical_slope = np.sqrt(2 * np.pi) * rho / a
    samplingtype = ['Langevin', 'Hamiltonian'] if sampling == 'both' else [sampling]
    # choose Wei and build simulate_trial partial
    plt.figure(figsize=(2.5,1.75))
    ax = plt.gca() 
    for type in samplingtype:
        if type == 'Langevin':
            Wei = 0
            label = 'Without SOM (Langevin)'
            ff_scale = 0.8* simulator.params['critical_weight']
        elif type == 'Hamiltonian':
            Wei = -0.6 * simulator.params['critical_weight']
            ff_scale = 1.8* simulator.params['critical_weight']
            label = 'With SOM (Hamiltonian)'
        else:
            raise ValueError(f"Invalid sampling type: {samplingtype}")
        simulate_trial = partial(
            simulator.run_simulation,
            Wei=Wei,
            ff_scale=ff_scale,
            Wee=simulator.params['recurrent_weight_e2e'],
            test_eq='normal'
        )

        # build an expanded list of Rf values (one entry per trial)
        expanded = [Rf for Rf in Rf_list for _ in range(n_trials)]

        # run all trials in parallel
        with mp.Pool(mp.cpu_count()) as pool:
            trial_results = list(tqdm(
                pool.imap(simulate_trial, expanded),
                total=len(expanded),
                desc=f"Running {n_trials} trials per Rf ({sampling})", file=sys.stdout
            ))

        # extract per-trial precision = 1/var(Se)
        precisions = np.array([
            1.0 / np.var(res['E_stim'][int(t_record/dt):])
            for res in trial_results
        ])
            # reshape into (n_Rf, n_trials) and average
        mean_prec = precisions.reshape(len(Rf_list), n_trials).mean(axis=1)

        plt.scatter(Rf_list,
                    mean_prec,
                    label=f'{label}Simulation (mean over {n_trials} trials)',
                    linewidth=2, alpha=0.7)
            # scatter of every trial
        # plt.scatter(expanded,
        #         precisions,
        #         s=20,
        #         alpha=0.6,
        #         label=f'{label} Simulation (all {int(len(expanded)/len(Rf_list))} trials)')

    # theoretical line
    plt.plot(Rf_list,
             theoretical_slope * np.array(Rf_list),
             label='Theoretical Prediction',
             linewidth=2, alpha=0.5, color='black')


    ax.set_xlim(left=0)
    plt.xlabel('Feedforward Intensity (Rf)')
    plt.ylabel('1 / Variance (Precision)')
    plt.title(f'Precision vs Input Intensity ({sampling}) — all trials')
    plt.legend()
    save_fig(plt.gcf(),
             f'precision_vs_Rf_mean_{sampling}({int(len(expanded)/len(Rf_list))}trials.eps',
             sampling)
    plt.close()


def plot_kl_divergence(param_dict, kl_dict, cmap, name='kl', T=False, tstart=0, sampling='Langevin'):
    """
    Plots the Kullback-Leibler divergence and normalized KL divergence of neural responses over time.

    Args:
    - param_dict: Dictionary containing simulation parameters
    - kl_dict: Dictionary containing KL divergences for different methods
    - cmap: Color map for plotting
    - name: Name for the plot (default is 'kl')
    - T: Simulation time (default is False)
    - tstart: Start time for plotting (default is 0)
    - sampling: Sampling method for saving the figure (default is 'Langevin')
    """
    if T == False:
        T = param_dict["simulation_time"]
    dt = param_dict["time_step"]
    t = np.arange(tstart, T+dt, dt)  
    # Plot KL divergence
    fig, ax2 = plt.subplots(figsize=(2.5, 1.75))
    for i, (method, kl) in enumerate(kl_dict.items()):
        if 'Natural Gradient' in method:
            ax2.semilogy(t, kl, label=method, color=cmap[int(i/2)], linestyle='--')
        else:
            ax2.semilogy(t, kl, label=method, color=cmap[int(i/2)])  # Changed to semilogy
    if tstart != 0:
        ax2.axvline(x=tstart, color='gray', linestyle='--', linewidth=1)
    ax2.set_ylabel('KL divergence (log scale)')
    ax2.set_xlim(left=0)
    ax2.set_title('KL divergence')
    ax2.set_xlabel(r'Time /$\tau$')
    ax2.legend()
    save_fig(fig, f'KL Divergence of {name}.eps', sampling=sampling)
    plt.close()
    # Plot normalized KL divergence
    fig, ax1 = plt.subplots(figsize=(2.5, 1.75))
    
    for i, (method, kl) in enumerate(kl_dict.items()):
        if 'Natural Gradient' in method:
            ax1.semilogy(t, kl/kl[0], label=method, color=cmap[int(i/2)], linestyle='--')
        else:
            ax1.semilogy(t, kl/kl[0], label=method, color=cmap[int(i/2)])  # Changed to semilogy
    if tstart != 0:
        ax1.axvline(x=tstart, color='gray', linestyle='--', linewidth=1)
    ax1.axvline(x=tstart, color='gray', linestyle='--', linewidth=1)
    ax1.set_ylabel('Normalized KL divergence (log scale)')
    ax1.set_title('Normalized') 
    ax1.set_xlabel(r'Time /$\tau$')
    ax1.set_xlim(left=0)
    ax1.legend()
    save_fig(fig, f'Normalized KL Divergence of {name}.eps', sampling=sampling)
    plt.close()
    


def plot_cross_correlation(cc_dict, cmap, name='cross_correlation', sampling='Langevin'):
    """
    Plots the cross-correlation of neural responses.

    Args:
    - cc_dict: Dictionary of cross-correlations for each method.
    - cmap: List of colors for plotting.
    - name: Name for the saved figure.
    - sampling: Label for figure saving context.
    """
    tLag = np.arange(0, 10.0, 0.01)
    fig, ax = plt.subplots(figsize=(5, 5))
    
    for i, (method, cc) in enumerate(cc_dict.items()):
        if 'Natural Gradient' in method:
            ax.plot(tLag, cc[:1000], label=method, color=cmap[int(i/2)], linestyle='--')
        else:
            ax.plot(tLag, cc[:1000], label=method, color=cmap[int(i/2)])
    ax.set_xlim(left=0)
    ax.set_xlabel(r'Time $\it{t}$ (/$\tau$)', size=16)
    ax.set_ylabel(r'Cross corr. $\rho_s$', size=16)
    ax.legend()
    save_fig(fig, f'Cross Correlation of {name}.eps', sampling=sampling)
    plt.close()


    
def Get_Kl_div_vs_Rf(params, Wee=0, num_trials=50, test_eq='eq',sampling='Langevin'):
    """
    Calculate Kullback-Leibler divergence and cross-correlation for different Rf values.
    
    Args:
    - params (dict): Dictionary containing simulation parameters.
    - Wee (int, optional): Excitatory-to-excitatory weight. Defaults to 0.
    - num_trials (int, optional): Number of simulation trials. Defaults to 50.
    - test_eq (str, optional): Type of test - 'eq' for equilibrium, 'non-eq' for non-equilibrium. Defaults to 'eq'.
    - sampling (str, optional): Type of sampling. Defaults to 'Langevin'.
    
    """
    
    simulator = CANNSimulator(params)
    simulator.initialize_network()
    
    kl_dict = {}
    se_dict = {}
    Ue_list = []
    cc_dict = {}
    Rf_list = np.linspace(1,5,3)

    if Wee ==0 :
        cmap = plt.cm.summer(np.linspace(0.3, 0.7, len(Rf_list)))  # Create color gradient
    else:
        cmap = plt.cm.summer(np.linspace(0.3, 0.7, len(Rf_list)))  # Create color gradient

    for Rf in tqdm(Rf_list, desc=f"Processing Rf at Wee={Wee}"):
        results = simulator.compute_bump_positions_height_over_trials(
            Rf, Wei=0, ff_scale=params['feedforward_scale'], Wee=Wee, num_trials=num_trials, test_eq=test_eq
        )
        
        Se_Lan_CANN = results['bump_positions'][:, int(params['t_steady'] / params['time_step'])-1:]
        Ue = results['bump_height']
        Ue_list.append(Ue)
        # kl_Lan_CANN = compute_kl_divergence(params, Se_Lan_CANN, Rf, num_trials=num_trials)
        cc_dict[f'CANN (Rf={Rf:.3f})'] = compute_cross_correlation(Se_Lan_CANN)
        kl_dict[f'CANN (Rf={Rf:.3f})'] = compute_kl_divergence(params, Se_Lan_CANN, Rf, num_trials=num_trials)
        # se_dict[f'CANN (Rf={Rf:.3f})'] = Se_Lan_CANN
        del Se_Lan_CANN, Ue
        gc.collect()

        
        if Wee ==0 :
            initials_states_list = results['bump_positions'][:,int(params['t_steady']/params['time_step'])-1]
            del results
            gc.collect()
            S_Lan_fisher = langevin_sampling(params, Rf, initials_states_list, num_trials=num_trials, test_eq=test_eq, conditioner=True)
            # kl_Lan_fisher = compute_kl_divergence(params, S_Lan_fisher, Rf, num_trials=num_trials)
            kl_dict[f'Natural Gradient (Rf={Rf:.3f})'] = compute_kl_divergence(params, S_Lan_fisher, Rf, num_trials=num_trials)
            # se_dict[f'Natural Gradient (Rf={Rf:.3f})'] = S_Lan_fisher
            cc_dict[f'Natural Gradient (Rf={Rf:.3f})'] = compute_cross_correlation(S_Lan_fisher)
            del S_Lan_fisher,initials_states_list
            gc.collect()
        else:
            del results
            gc.collect()               
    if test_eq == 'eq':
        name=f"Different Rf at equilibrium at Wee={Wee} at {params['initial_var_eq']:.3f} {params['simulation_time']:.3f}"
    elif test_eq == 'non-eq':
        name=f"Different Rf at non-equilibrium for Wee={Wee} at {params['initial_var_eq']:.3f} {params['simulation_time']:.3f}"
    plot_kl_divergence(params, kl_dict, cmap = cmap, name = name, T=params['simulation_time'] - params['t_steady'],sampling = 'Langevin')

    plot_cross_correlation(cc_dict, cmap, name=name, sampling=sampling)
    

    dt = params['time_step']
    t_steady = params['t_steady']
    plt.figure()
    for i, Rf in enumerate(Rf_list):
        plt.plot(np.arange(0,t_steady,dt), np.average(Ue_list[i],axis=0)[:int(t_steady/dt)], 
                label=f'Bump height for RF = {Rf}', color=cmap[i])
        fisher = Rf * simulator.params['feedforward_scale']
        plt.plot(np.arange(0,t_steady,dt), fisher * np.ones(int(t_steady/dt)), 
            label=f'$\tau$ FI for RF = {Rf}', linestyle='--', color=cmap[-1])
    ax = plt.gca()  # Create the ax object
    ax.set_xlim(left=0)
    plt.xlabel(r'Time /$\tau$') 
    plt.ylabel('Bump height')
    plt.legend()   
    save_fig(plt.gcf(),f'Bump height for {name}.eps',sampling=sampling)
    plt.close()


    


def Get_Kl_div_vs_Wee(params,Wee_list = np.linspace(0, 1, 3),Rf = 3, num_trials=50,test_eq = 'eq',sampling = 'Langevin'):
    """
    Calculate Kullback-Leibler divergence and cross-correlation for different wee values.
    
    Args:
    - params (dict): Dictionary containing simulation parameters.
    - Wee_list (list /array , optional): List of wee values. Defaults to np.linspace(0, 1, 3).
    - Rf (float, optional): Input intensity for Langevin sampling. Defaults to 3.
    - num_trials (int, optional): Number of simulation trials. Defaults to 50.
    - test_eq (str, optional): Type of test - 'eq' for equilibrium, 'non-eq' for non-equilibrium. Defaults to 'eq'.
    - sampling (str, optional): Type of sampling. Defaults to 'Langevin'.
    
    """
    simulator = CANNSimulator(params)
    simulator.initialize_network()
    # simulator.params['feedforward_intensity_scale'] * 0.8 
    kl_dict = {}
    se_dict = {}
    cc_dict = {}
    Ue_list = []


    cmap = plt.cm.autumn(np.linspace(0.3, 0.7, len(Wee_list)))  # Create color gradient

    for Wee in tqdm(Wee_list, desc="Processing Wee"):
        print(Wee)
        results = simulator.compute_bump_positions_height_over_trials(Rf, Wei = 0, ff_scale = params['feedforward_scale'], Wee = Wee,num_trials=num_trials,test_eq=test_eq)        
        Se_Lan_CANN = results['bump_positions'][:,int(params['t_steady']/params['time_step'])-1:]
        if Wee ==0 :
            initials_states_list = results['bump_positions'][:,int(params['t_steady']/params['time_step'])-1]

        Ue = results['bump_height']
        Ue_list.append(Ue)
        kl_Lan_CANN = compute_kl_divergence(params,Se_Lan_CANN,Rf,num_trials = num_trials)
        kl_dict[f'CANN (Wee={Wee:.3f})'] = compute_kl_divergence(params,Se_Lan_CANN,Rf,num_trials = num_trials)
        cc_dict[f'CANN (Wee={Wee:.3f})'] = compute_cross_correlation(Se_Lan_CANN)

        # se_dict[f'CANN (Wee={Wee:.3f})'] = Se_Lan_CANN
  
        
    S_Lan_fisher = langevin_sampling(params, Rf, initials_states_list, test_eq=test_eq, conditioner=True, num_trials=num_trials)

    kl_dict['Natural Gradient'] = compute_kl_divergence(params, S_Lan_fisher, Rf)
    # se_dict['Natural Gradient'] = S_Lan_fisher
    cc_dict['Natural Gradient'] = compute_cross_correlation(S_Lan_fisher)

    if test_eq == 'eq':
        name = 'Different Wee at equilibrium for Rf = {}'.format(Rf)
    elif test_eq == 'non-eq':

        name = 'Different Wee at non-equilibrium for Rf = {}'.format(Rf)
    else:
        print(test_eq)
        name = 'Different Wee at non-equilibrium for Rf = {}'.format(Rf)
    if test_eq == 'non-eq':
        plot_kl_divergence(params, kl_dict, cmap = cmap, name = name, T=params['simulation_time'],tstart=params['t_steady'],sampling = 'Langevin')

    plot_cross_correlation(cc_dict, cmap, name=name, sampling=sampling)
    plt.close()
    dt = params['time_step']
    # t_steady = params['t_steady']
    T= params['simulation_time']
    plt.figure()
    for i, Wee in enumerate(Wee_list):
        plt.plot(np.arange(0,T,dt), np.average(Ue_list[i],axis=0), 
                label=f'Bump height for Wee = {Wee}', color=cmap[i])
    fisher = Rf * simulator.params['feedforward_scale']
    plt.plot(np.arange(0,T,dt), fisher * np.ones(int(T/dt)), 
        label=r'$\tau$ FI', linestyle='--', color=cmap[0])
    ax = plt.gca()  # Create the ax object
    ax.set_xlim(left=0)
    plt.xlabel(r'Time /$\tau$') 
    plt.ylabel('Bump height')
    plt.legend()   
    save_fig(plt.gcf(),f'Bump height for different wee{name}.eps',sampling=sampling)
    plt.close()


def plot_bump_height_vs_time_const(params, Wee_list,Rf =2 , num_trials=50,test_eq="normal"):
    """
    Plot the relationship between bump height and time constant for given parameters.

    Args:
    - params (dict): Dictionary of simulation parameters.
    - Wee_list (list): List of excitatory-to-excitatory connection weights.
    - Rf (int, optional): input intensity. Default is 2.
    - num_trials (int, optional): Number of simulation trials. Default is 50.
    - test_eq (str, optional): Type of test equation. Default is "normal".
    """
    
    t_steady = params['t_steady']
    recording_start = params['recording_start']
    dt = params['time_step']
    simulator = CANNSimulator(params)
    simulator.initialize_network()
    # simulator.params['feedforward_intensity_scale'] * 0.8 
    Ue_list = []
    tc_list = []
    for Wee in tqdm(Wee_list, desc="Processing Wee"):
        results = simulator.compute_bump_positions_height_over_trials(Rf, Wei = 0, ff_scale = params['feedforward_scale'], Wee = Wee,num_trials=num_trials,test_eq=test_eq)        
        Se_Lan_CANN = results['bump_positions']#[:,int(params['t_steady']/params['time_step'])-1:]
        Ue = np.average(np.average(results['bump_height'][:,int(recording_start/dt)-1:],axis=0)) #
        Ue_list.append(Ue)
        cc = compute_cross_correlation(Se_Lan_CANN,int(recording_start/dt))
        timeconst = extract_time_constants(cc)*params['time_step']
        tc_list.append(timeconst)
    plt.figure(figsize=(2.5,1.75))
    ax = plt.gca()  # Create the ax object
    np.save('Ue_list.npy', Ue_list)
    np.save('tc_list.npy', tc_list)
    plt.scatter(Ue_list,tc_list)
    tau =  params['time_constant_exc']
    rho = simulator.params['neuron_density']
    a = simulator.params['gaussian_width_exc']
    ff = simulator.params['feedforward_scale']
    plt.plot(Ue_list,np.array(Ue_list) *tau/(ff*Rf), color='black')
    ax.set_xlim(left=0)
    ax.set_ylim(bottom=0)
    plt.xlabel('Bump height')
    plt.ylabel('Time constant')
    plt.title(f'Time constant vs Bump height')
    save_fig(plt.gcf(), f'Time constant vs Bump height2.eps',sampling = 'Langevin')
    

def plot_bump_height_vs_Rf(params, Wee_list, Rf_list, test_eq="normal", noise=False):
    """
    Plot the relationship between bump height and Fisher Information for given parameters.

    Args:
    - params (dict): Dictionary of simulation parameters.
    - Wee_list (list/array): List of excitatory-to-excitatory connection weights.
    - Rf_list (list/array): List of input intensities.
    - test_eq (str, optional): Type of test equation. Default is "normal".
    - noise (bool, optional): Flag indicating whether noise is included in the simulation. Default is False.
    """
    a = params["gaussian_width_exc"]
    simulator = CANNSimulator(params)
    simulator.initialize_network()
    rho = simulator.params['neuron_density']
    Ue_dict = {}
    for Wee in Wee_list:
        Ue_list =[]
        for Rf in tqdm(Rf_list, desc="Processing Rf"):
            results = simulator.run_simulation(Rf, Wei=0, ff_scale=params['feedforward_scale'], Wee=Wee, test_eq=test_eq, noise=noise)        

            Ue = np.average(results['E_bump_height'][int(params['t_steady']*2/params['time_step']):])
            Ue_list.append(Ue)
        print(Ue_list)
        Ue_dict[Wee] = Ue_list
    cmap = plt.cm.autumn(np.linspace(0.3, 0.7, len(Wee_list)+1))  # Create color gradient
    plt.figure(figsize=(2.5, 1.75))
    ax = plt.gca()  # Create the ax object
    
    for i, Wee in enumerate(Wee_list):
        plt.plot((np.sqrt(2 * np.pi) * rho / a) * Rf_list, Ue_dict[Wee], label=f'Wee = {Wee:.3f}', color=cmap[i])
    plt.xlabel('Fisher Information')
    plt.ylabel('Bump height')
    ax.set_xlim(left=0)
    plt.title(f'Bump height vs Fisher Information')
    plt.legend()
    save_fig(plt.gcf(), f'Bump height vs Fisher Information.eps', sampling='Langevin')
    for i, Wee in enumerate(Wee_list):
        plt.plot(Rf_list, Ue_dict[Wee], label=f'Wee = {Wee:.3f}', color=cmap[i])
    plt.xlabel('Fisher Information')
    plt.ylabel('Bump height')
    ax.set_xlim(left=0)
    plt.title(f'Bump height vs Fisher Information')
    plt.legend()
    save_fig(plt.gcf(), f'Bump height vs Input Intensity.eps', sampling='Langevin')


def plot_conv_vs_Rf(params, Wee_list, Rf_list, num_trials=50, test_eq="eq",dir = "saved"):
    """
    runs simulations for each Wee, collects convergence times,
    and plots Convergence Time vs Rf.
    """
    # 1) initialize simulator once
    simulator = CANNSimulator(params)
    simulator.initialize_network()
    
    # 2) collect data
    all_conv = []
    fisher_conv = []
    for Wee in Wee_list:
        conv, conv_f = compute_conv_times_for_Wee(
            params, simulator, Wee, Rf_list, num_trials, test_eq, save_dir=dir
        )

        all_conv.append(conv)
        # only fill once (Wee==0)
        if Wee == Wee_list[0]:
            fisher_conv = conv_f
    
    # 3) plotting
    cmap = plt.cm.autumn(np.linspace(0.3, 0.7, len(Wee_list)+1))
    plt.figure(figsize=(2.5,1.75))
    for i, Wee in enumerate(Wee_list):
        print(Wee)
        plt.plot(Rf_list, all_conv[i]*params['time_step'],
                 label=f'Wee = {Wee}',
                 color=cmap[i])
    # if fisher_conv.any():
    plt.plot(Rf_list, fisher_conv*params['time_step'],label = 'Natural Gradient', color=cmap[-1], linestyle='--')
    plt.xlabel('Feedforward Intensity (Rf)')
    plt.ylabel(r'Convergence Time \ $\tau$')
    plt.title('Convergence Time vs Feedforward Intensity')
    plt.legend()
    plt.xlim(left=0)
    
    save_fig(plt.gcf(),
             'Convergence time vs Feedforward Intensity2.eps',
             sampling='Langevin')
    plt.close()


# Updated plotting function to load saved bumps and plot convergence times
def plot_conv_vs_Rffromdic(params, Wee_list, Rf_list, num_trials=50, test_eq="eq", data_dir="saved", load_saved=False):
    """
    Plots Convergence Time vs Rf for given Wee_list.
    If load_saved=True, loads precomputed 'bumps' arrays from data_dir and computes convergence times.
    Otherwise, runs simulations and saves bumps as before.
    """
    # 1) initialize simulator if needed
    simulator = CANNSimulator(params) if not load_saved else None
    if simulator:
        simulator.initialize_network()

    # 2) collect data
    all_conv = []
    fisher_conv = []
    for idx, Wee in enumerate(Wee_list):
        if load_saved:
            # load saved bumps, compute conv times
            conv = []
            conv_f = []
            for Rf in Rf_list:
                file_path = os.path.join(data_dir, f"bumps_Wee{Wee:.3f}_Rf{Rf:.3f}_nt{num_trials}.npy")
                bumps = np.load(file_path)
                kl = compute_kl_divergence(params, bumps, Rf, num_trials=num_trials)
                conv.append(get_convergence_time(kl))

                if Wee == Wee_list[0]:
                    # fisher variant load, assuming saved as 'fisher_Wee0_Rf*.npy'
                    fisher_file = os.path.join(data_dir, f"samp_fisher_Rf{Rf:.3f}_nt{num_trials}.npy")
                    if os.path.exists(fisher_file):
                        samp = np.load(fisher_file)
                        klf = compute_kl_divergence(params, samp, Rf, num_trials=num_trials)
                        conv_f.append(get_convergence_time(klf))
                    else:
                        initials = bumps[:, 0]
                        samp = langevin_sampling(
                            params, Rf, initials, num_trials=num_trials,
                            test_eq=test_eq, conditioner=True
                        )
                        klf = compute_kl_divergence(params, samp, Rf, num_trials=num_trials)
                        conv_f.append(get_convergence_time(klf))
                        fisher_file = os.path.join(data_dir, f"samp_fisher_Rf{Rf:.3f}_nt{num_trials}.npy")
                        np.save(fisher_file, samp)
            all_conv.append(np.array(conv))
            if Wee == Wee_list[0]:
                fisher_conv = np.array(conv_f)
        else:
            # compute and save as before
            conv, conv_f = compute_conv_times_for_Wee(
                params, simulator, Wee, Rf_list, num_trials, test_eq, save_dir=data_dir
            )
            all_conv.append(conv)
            if Wee == Wee_list[0]:
                fisher_conv = conv_f

    np.save(os.path.join(data_dir, f"all_conv_Wee{Wee_list}.npy"), all_conv)
    np.save(os.path.join(data_dir, f"fisher_conv_Wee{Wee_list}.npy"), fisher_conv)
    # 3) plotting
    cmap = plt.cm.autumn(np.linspace(0.3, 0.7, len(Wee_list)+1))
    plt.figure(figsize=(2.5,1.75))
    
    for i, Wee in enumerate(Wee_list):
        plt.plot(Rf_list, all_conv[i] * params['time_step'],
                 label=f'Wee = {Wee}', color=cmap[i])
    if load_saved and fisher_conv.size:
        plt.plot(Rf_list, fisher_conv * params['time_step'],
                 label='Natural Gradient', color=cmap[-1], linestyle='--')
    elif not load_saved and len(fisher_conv):
        plt.plot(Rf_list, fisher_conv * params['time_step'],
                 label='Natural Gradient', color=cmap[-1], linestyle='--')

    plt.xlabel('Feedforward Intensity (Rf)')
    plt.ylabel(r'Convergence Time \ $\tau$')
    plt.title('Convergence Time vs Feedforward Intensity')
    plt.legend()
    plt.xlim(left=0)

    save_fig(plt.gcf(), 'Convergence_time_vs_Feedforward_Intensity.eps', sampling='Langevin')
    plt.close()


def plot_possion_ff_likelihood(params,I):
    """
    Draws the two-panel figure:
      1) Population response: Poisson spike counts vs. neuron index θ_j
      2) Gaussian likelihood approximation of p(I^f | s)
    """

    # N = params['num_neurons']
    mu  = params['input_position']
    a = params['gaussian_width_exc']
    # Preferred angles and firing rates
    theta_j = np.linspace(-180, 180, 31)
    lambda_j = I * np.exp(- (theta_j - mu)**2 / (2 * a**2))
    
    # Sample Poisson counts
    counts = np.random.poisson(lambda_j)
    
    # Population vector estimate and precision
    n_r = counts.sum()
    x_hat = np.sum(counts * theta_j) / n_r
    Lambda = n_r / (a**2)
    
    # 1) Population Response
    plt.figure(figsize=(2.5,1.75))
    plt.bar(theta_j, counts)
    # plt.axhline(y=x_hat, color='black', linestyle='-')
    # plt.annotate(
    #     'x (observed feature)',
    #     xy=(x_hat, max(counts)*0.6), xytext=(x_hat + 30, max(counts)*0.8),
    #     arrowprops=dict(arrowstyle='->')
    # )
    plt.gca().set_xlim(-180, 180)
    plt.gca().set_xticks([-180,0,180])
    plt.xlabel('Neuron index θ (°)')
    plt.ylabel('Feedforward input $I^{(f)}$')
    plt.title('Population Response')
    save_fig(plt.gcf(), f'Population Response.eps', sampling='Langevin')
    
    # 2) Gaussian Likelihood Approximation
    s_range = np.linspace(x_hat - 50, x_hat + 50, 400)
    likelihood = np.exp(- Lambda * (s_range - x_hat)**2 / 2)
    likelihood /= np.max(likelihood)
    
    plt.figure(figsize=(2.5,1.75))
    plt.plot(s_range, likelihood)
    plt.axvline(x=x_hat, linestyle='-')
    plt.annotate('x', xy=(x_hat, 1.0), xytext=(x_hat + 5, 0.8),
                 arrowprops=dict(arrowstyle='->'))
    plt.text(x_hat, 0.5, f'Λ = {Lambda:.1f}', ha='center')
    plt.xlabel('Stimulus s')
    plt.ylabel('Likelihood $p(I^{(f)}\\mid s)$')
    plt.title('Gaussian Likelihood Approximation')
    save_fig(plt.gcf(), f'Gaussian Likelihood Approximation.eps', sampling='Langevin')

def plot_one_attractor(params, Rf=10,test_eq="normal",tstart = 0,tend = 100):
    """
    Plot one simulation for Langevin sampling

    Args:
    - params: dictionary containing simulation parameters
    - Rf: float, input intensity for Langevin sampling
    - test_eq: str, type of input used for initialization
        -eq: get equilibrium results
        -non-eq: get non-equilibrium results
        -nomal: get normal results/ the initialization is normal
    - tstart: int, start time for plotting
    - tend: int, end time for plotting
    """
    
    simulator = CANNSimulator(params)
    simulator.initialize_network()
    results_Lan = simulator.run_simulation(
        Rf = Rf,#0.8 * simulator.params['feedforward_intensity_scale'], 
        Wei= 0, 
        ff_scale=params['feedforward_scale'],
        Wee = simulator.params['recurrent_weight_e2e'],
        test_eq= test_eq
    )
    dt = params['time_step']
    recording_start = params['recording_start']
    fr = results_Lan['E_FR'][:,int(recording_start/dt):int(tend/dt)]
    plt.figure(figsize=(6,6))
    ax = plt.gca()  # Create the ax object

    plt.plot(simulator.params['PrefStim'],np.mean(fr,axis=1), color='blue')
    plt.fill_between(simulator.params['PrefStim'],np.mean(fr,axis=1)-np.std(fr,axis=1),np.mean(fr,axis=1)+np.std(fr,axis=1),color='lightsteelblue')
    ax.set_xticks([-180,-120,-60,0,60,120,180])
    plt.xlabel('Preferred Stimulus')
    plt.ylabel('Firing Rate')
    plt.title('Firing Rate vs Preferred Stimulus')
    save_fig(plt.gcf(), f'Firing Rate vs Preferred Stimulus.eps', sampling='Langevin')
    plt.close()
    plt.figure(figsize=(6,6))
    ax = plt.gca()  # Create the ax object
    si = results_Lan['Synaptic_Input'][:,int(recording_start/dt):int(tend/dt)]
    plt.plot(simulator.params['PrefStim'],np.mean(si,axis=1), color='blue')
    plt.fill_between(simulator.params['PrefStim'],np.mean(si,axis=1)-np.std(si,axis=1),np.mean(si,axis=1)+np.std(si,axis=1),color='lightsteelblue')
    ax.set_xticks([-180,-120,-60,0,60,120,180])
    plt.xlabel('Preferred Stimulus')
    plt.ylabel('Synaptic Input')
    plt.title('Synaptic Input vs Preferred Stimulus')
    save_fig(plt.gcf(), f'Synaptic Input vs Preferred Stimulus.eps', sampling='Langevin')
    plt.close()

def plot_jsd_heatmap_with_min_line(csv_path, critical_weight, rho = 0.5,markers=None, save_path=None):
    """
    Plot JS divergence heatmap from a CSV file, overlay the min-JSD path,
    and invert the y-axis.

    Args:
        csv_path (str): Path to the CSV file containing columns: Wei, Wef, JSD.
        critical_weight (float): The critical weight for normalization.
        markers (list of tuples): List of (Wef, Wei, color) tuples for dot markers.
        save_path (str or None): If provided, saves the plot to this path instead of showing.
                                 (Note: current code saves based on csv_path name)
    """
    # Load data
    df = pd.read_csv(csv_path)

    # Normalize axes to show as multiples of critical_weight
    df['Wei_unit'] = df['Wei'] / critical_weight
    df['FF_unit'] = df['Wef'] / critical_weight

    # Create pivot table for heatmap
    # Ensure Wei_unit is sorted ascendingly for consistent y-axis indexing later
    heatmap_data = df.pivot(index='Wei_unit', columns='FF_unit', values='JSD')
    heatmap_data = heatmap_data.sort_index(ascending=True)  # Sort Wei_unit (index) ascending

    # Prepare min-JSD line data
    min_wei_for_each_ff = heatmap_data.idxmin(axis=0)  # Find min-JSD row (Wei_unit) for each FF_unit
    ff_actual_vals = min_wei_for_each_ff.index.values  # Actual FF_unit values
    wei_actual_vals = min_wei_for_each_ff.values       # Corresponding actual Wei_unit values

    # Start plot
    plt.figure(figsize=(6, 6))
    ax = plt.gca()  # Create the ax object
    plt.imshow(
        heatmap_data,
        cmap="Blues", 
        vmin=0, vmax=0.3
    )
    plt.colorbar(label='JS Div.')
    # Set X-axis ticks and labels (using actual FF_unit values for labels)
    # --- X-axis Ticks Modification ---
    desired_x_labels_str = np.arange(0, 1.6, 0.2).astype(str)
    desired_x_values_num = [float(s)*(2*np.sqrt(2)) for s in desired_x_labels_str]
    
    ff_unit_axis_values = heatmap_data.columns.to_numpy()
    axis_min_ff_unit = ff_unit_axis_values.min()
    axis_max_ff_unit = ff_unit_axis_values.max()

    tick_infos = {} # Using dict to store {index: (label_str, distance_to_target_val)}
                    # This helps select the best label if multiple targets map to the same index.

    for target_val, target_label_str in zip(desired_x_values_num, desired_x_labels_str):
        # Only consider target values that are reasonably within the data's FF_unit range
        # Add a small tolerance (e.g., 0.05) for floating point comparisons at the edges
        if (target_val >= axis_min_ff_unit - 0.05) and (target_val <= axis_max_ff_unit + 0.05):
            idx = np.abs(ff_unit_axis_values - target_val).argmin()
            distance = np.abs(ff_unit_axis_values[idx] - target_val)

            # If this index is not taken, or if this target_val is a closer match for this index
            if idx not in tick_infos or distance < tick_infos[idx][1]:
                tick_infos[idx] = (target_label_str, distance)
            
    if tick_infos:
        sorted_tick_indices = sorted(tick_infos.keys())
        final_x_tick_positions = sorted_tick_indices
        final_x_tick_labels = [tick_infos[idx][0] for idx in sorted_tick_indices]
        
        ax.set_xticks(final_x_tick_positions)
        ax.set_xticklabels(final_x_tick_labels, rotation=45, ha="right")
    else:
        # Fallback to default ticks if no desired ticks are found in range
        num_default_xticks = 6 # Or 5 as per previous y-axis
        default_x_indices = np.linspace(0, len(heatmap_data.columns) - 1, num_default_xticks, dtype=int, endpoint=True)
        ax.set_xticks(default_x_indices)
        ax.set_xticklabels([f"{heatmap_data.columns[i]:.1f}" for i in default_x_indices], rotation=45, ha="right")
    # --- End X-axis Ticks Modification ---lumns[i]:.1f}" for i in [1, 10, 20, 30, 40]])
    # ax.set_xticklabels([f"{heatmap_data.columns[i]:.2f}" for i in x_ticks], rotation=45, ha="right")

    # Set Y-axis ticks and labels (using actual Wei_unit values for labels)
    y_ticks = np.linspace(0, len(heatmap_data.index) - 1, num=5, dtype=int)
    ax.set_yticks(y_ticks)
    ax.set_yticklabels([f"{heatmap_data.index[i]:.2f}" for i in y_ticks])

    # Invert Y-axis: Smallest Wei_unit values will be at the bottom
    ax.invert_yaxis()

    # Axis labels
    plt.xlabel(r'Feedfwd. $w_{EF}$ $(/w_C)$')
    plt.ylabel(r'Inh. $w_{ES}$ $(/w_C)$')
    plt.title('JSD Heatmap with Min-JSD Path') # Added a title for clarity

    # Plot yellow minimum-JSD path using cell indices
    # x_idxs are the integer indices of the columns [0, 1, 2, ...]
    x_idxs = np.arange(len(heatmap_data.columns))

    # y_idxs are the integer indices of the rows corresponding to min JSD Wei_unit values
    # We use .get_loc() on the sorted heatmap_data.index
    y_idxs = [heatmap_data.index.get_loc(w) for w in wei_actual_vals]
    
    plt.plot(x_idxs, y_idxs, color='yellow', linewidth=2, zorder=10, label='Min JSD Path')
    if any(y_idxs): # Add legend if there's a line to label
        plt.legend()


    # Optional markers: these use cell indices as well
    if markers:
        for ff, wei, color in markers:
            # Find the closest column index for ff (actual FF_unit value)
            x_idx = np.abs(heatmap_data.columns.to_numpy() - ff/critical_weight).argmin()
            # Find the closest row index for wei (actual Wei_unit value)
            y_idx = np.abs(heatmap_data.index.to_numpy() - wei/critical_weight).argmin()
            # x_idx = [heatmap_data.columns.get_loc(ff_actual_vals[y_idx])]
            plt.scatter(x_idx, y_idx, color=color, edgecolors='black', s=100, zorder=11)
        # if markers: # Add legend if there are markers
        #      ax.legend(bbox_to_anchor=(1.05, 1), loc='upper left')


    plt.tight_layout()

    # Save the plot
    base_name = os.path.splitext(os.path.basename(csv_path))[0]
    png_name = base_name + ".png"
    eps_name = base_name + ".eps"

    if save_path:
        # If a specific save_path directory is given
        if not os.path.isdir(save_path):
            os.makedirs(save_path, exist_ok=True)
        png_name = os.path.join(save_path, png_name)
        eps_name = os.path.join(save_path, eps_name)

    plt.savefig(png_name)
    print(f"Plot saved to {png_name}")
    
    # Using the custom save_fig for EPS
    save_fig(plt.gcf(), eps_name, sampling='Langevin') # Removed sampling if not standard

    if save_path is None : #Only show if not saving to a specific path in args (still saves locally)
        plt.show()
    plt.close()

def plot_tuning_curves(n_neurons=9, variance=40, sigma=40):

    z = np.linspace(-180, 180, 1000)
    preferred_stimuli = np.linspace(-180, 180, n_neurons)

    plt.figure(figsize=(6, 4))
    cmap = get_cmap('Purples')
    for i, mu in enumerate(preferred_stimuli):
        color = cmap(i / (n_neurons - 1))  # Normalize index to [0, 1]
        firing_rate = np.exp(-0.5 * ((z - mu) / sigma)**2)
        plt.plot(z, firing_rate, color=color)
    ax = plt.gca()
    plt.xlabel('Stimulus feature $z$')
    plt.ylabel('Firing rate')
    plt.xlim([-180, 180])
    plt.ylim([0, 1.1])
    ax.set_xticks([-180, -120,-60,0,60,120, 180])
    # ax.set_xticks([-180, -120,-60,0,60,120, 180])
    plt.tight_layout()
    save_fig(plt.gcf(), 'Tuning Curves.eps', sampling='Langevin')
```
