## Supplementary for "Natural gradient Bayesian sampling automatically emerges in canonical cortical circuits": 2d.py

```
from cannND import *
from analysis_tools2D import *
from plotdataND import *
import warnings
warnings.filterwarnings('ignore')

params = {
            'time_constant_exc': 1.0,
            'position_max': 180.0,
            'position_min': -180.0,
            'gaussian_width_exc': 40.0,
            'gaussian_width_ES': 20.0,
            'num_neurons': 180,
            'simulation_time': 500.0,
            'time_step': 0.01,
            'recording_start': 50,
            'Fano_factor': 0.5,
            'normalization_k': 0.0005,
            'inhibitory_gain': 10,
            'input_position': [-2,2],
            'feedforward_scale': 0.71648,
            't_steady':50,
            'initial_mean_eq':0,
            'initial_var_eq':30, 
            'Dimension': 2
        }

if __name__ == "__main__":

    # Show the simulation result for one run CANN model
    #Langevin simulation
    #Figure 3 B C 2d population, the bump postition read out
    plot_one_simulationND(params = params, Rf_both = [5,10] ,normal_input =True,tstart = 50, tend=60)
    #Figure 3D vector field of cann and the heatmap of posterior and trajectory  
    plot_bump_position2D(params, Rf_both = [5,10] ,sampling="Langevin",normal_input =True)

    
    #Figure 3E 
    simulator = CANNSimulator2D(params)  # Changed to 2D simulator
    simulator.initialize_network()
    Rf_both = [5,10]
    Lambda_s_opt, KLD = simulator.find_prior_precision(Wee=0, Rf_both=Rf_both,normal_input=True)
    plot_vector_field_LAN(params,Rf_both = Rf_both, Wei=0, ff_scale = params['feedforward_scale'],Wee = 0, Lambda_s= Lambda_s_opt,fisher = "both")
    
    # Figure 3F
    num_trials=500
    simulator = CANNSimulator2D(params)  # Changed to 2D simulator
    simulator.initialize_network()
    Rf_both = [5,10]
    Wee = 0
    # if Rf, wee are changed, re-run the find_prior_precision function to get the new Lambda_s_opt
    Lambda_s_opt, KLD = simulator.find_prior_precision(Wee=0, Rf_both=Rf_both,normal_input=True)
    Get_Kl_div_vs_Lan_eq2D(params,Wee = Wee, Rf_both = Rf_both,  num_trials=num_trials, normal_input=False, Lambda_s = Lambda_s_opt)
    
    # SI fig2 
    plot_precision_prior(params,num_trials=50)
    # if you have the precision prior data, you can load it directly 
    # plot_precision_prior_fromdic(filename = 'precision_prior2.npy')
    plot_diagonal_prior(lambda_s=0.0004)
    plot_prior_L_vs_wcoup(params)
```
