## Supplementary for "Natural gradient Bayesian sampling automatically emerges in canonical cortical circuits": analysis_tools2D.py

```
import numpy as np
from tqdm import tqdm
from sklearn.linear_model import LinearRegression
from cannND import CANNSimulator2D
import multiprocessing as mp
from functools import partial
from scipy.linalg import sqrtm
# from numba import njit
# @njit
def compute_posterior_precision(Rf_both, param_dict,Lambda_s=None):
    """
    Computes the prior precision matrix for a 2D CANN.

    Args:
        Rf_both (tuple): (Rf1, Rf2) scaling factors for the likelihood.
        param_dict (dict): Network parameters, must include 'input_position'.
        Lambda_s (float, optional): Prior precision; if None, it will be computed.

    Returns:
        invCovPost (ndarray): Inverse covariance matrix of the prior precision.
        muPost (ndarray): Mean of the prior precision.
    """
    # Extract key parameters
    D = param_dict["Dimension"]
    rho = param_dict['num_neurons'] / (param_dict['position_max'] - param_dict['position_min'])
    a = param_dict['gaussian_width_exc']

    # Likelihood precision
    Rf1, Rf2 = Rf_both
    invCovLH = (np.sqrt(2 * np.pi) * rho) / a * np.diag([Rf1, Rf2])

    # Prior precision matrix (structured for 2D)
    mu = param_dict['input_position']
    

    prior_precision = 2 * np.eye(D) - np.ones((D, D))
    invCovPost = Lambda_s * prior_precision + invCovLH
    muPost = np.linalg.solve(invCovPost, invCovLH @ mu)

    return invCovPost, muPost
def compute_kl_divergence2D(param_dict,Lambda_s,Se1,Se2,Rf_both,num_trials = 50):
    """
    Computes the Kullback-Leibler divergence between sampling from CANN and reference distributions from feedforward input.    
    Returns:
        kl_all: A list of tuples containing the computed KLD values over time for each trial.
    """

    # Load parameters
    D = param_dict["Dimension"]
    rho = param_dict['num_neurons'] / (param_dict['position_max'] - param_dict['position_min'])
    a = param_dict['gaussian_width_exc']
    
    Rf1,Rf2 = Rf_both
    invCovLH = (np.sqrt(2 * np.pi) * rho) / a *np.eye(2)
    invCovLH[0,0] = invCovLH[0,0] * Rf1
    invCovLH[1,1] = invCovLH[1,1] * Rf2
    mu = param_dict['input_position']
    if isinstance(mu,int):
        mu = np.array([mu, -mu])
    muLH = mu
    prior_precision_structure = 2 * np.eye(D) - np.ones((D, D))
    invCovPost = Lambda_s * prior_precision_structure + invCovLH
    muPost = np.linalg.solve(invCovPost, invCovLH @ muLH)
    # The covariance of the posterior is the inverse of the precision matrix.
    if np.linalg.det(invCovPost) <= 1e-6:
        raise ValueError("Determinant of the inverse covariance matrix is too small.")
    covPost = np.linalg.inv(invCovPost)

    counter = np.arange(1, Se1.shape[1]+1, 1)
    kl_all = []

    for i in tqdm(counter, desc="Computing 2D KL divergence"):
        Sshort1 = Se1[:, :i].reshape(-1)
        Sshort2 = Se2[:, :i].reshape(-1)
        SeMean = [np.mean(Sshort1), np.mean(Sshort2)]
        SeCov = np.cov(Sshort1, Sshort2)
        kl = get_KL_div(SeMean, SeCov, muPost, covPost)
        kl_all.append(kl)

    return kl_all

def get_KL_div(mu1, cov1, mu2, cov2):
    """
    Compute the KL divergence between two multivariate Gaussian distributions.
    Computes KL(P||Q) where P ~ N(mu1, cov1), Q ~ N(mu2, cov2)
    """
    d = len(mu1)
    inv_cov2 = np.linalg.inv(cov2)
    diff = mu2 - mu1
    term1 = np.trace(inv_cov2 @ cov1)
    term2 = diff.T @ inv_cov2 @ diff
    term3 = np.log(np.linalg.det(cov2) / np.linalg.det(cov1))
    return 0.5 * (term1 + term2 - d + term3)

def Lan_run_trial2D(T, dt, rate, precision, mean, initial_state):
    """Function must be at the top level for pickling.
       initial_state should be a tuple or array-like with two values."""
    num_steps = int(T/dt) + 1
    trial = np.zeros((num_steps, 2))
    trial[0, :] = initial_state  # Unpack the tuple/array

  #  noise = np.random.normal(0, 1, (num_steps - 1, 2))
    
    sqrtm_result = sqrtm(2 * rate * dt)
    # if sqrtm_result.shape[0] != sqrtm_result.shape[1] or sqrtm_result.shape[0] != noise[0, :].shape[0]:
    #     raise ValueError("Dimension mismatch: sqrtm result must be square and match the dimensions of noise.")
    
    for t in range(1, num_steps):
        drift_term = rate @ precision @ (-trial[t-1, :] + mean) * dt
        noise_term = sqrtm_result @ np.random.normal(0, 1, (2,))
        trial[t, :] = trial[t-1, :] + drift_term + noise_term
    
    return trial


def langevin_sampling2D(param_dict, Rf_both,Lambda_s, initial_states_list, num_trials=50, normal_input=False, Diag=True):
    """
    Perform Langevin sampling in 2D space using Natural Gradient.

    Args:
    - param_dict (dict): Dictionary containing simulation parameters.
    - Rf_both (tuple): Tuple containing scaling factors for Rf1 and Rf2.
    - Lambda_s (float): Lambda parameter for the posterior precision.
    - initial_states_list (list): List of initial states for each trial.
    - num_trials (int): Number of trials to run (default is 50).
    - normal_input (bool): Flag indicating whether to use normal input (default is False).
    - Diag (bool): Flag indicating whether to use diagonal Fisher information matrix (default is True).

    Returns:
    - results (np.array): Array of results from the Langevin sampling trials.
    """

    T = param_dict["simulation_time"] - param_dict['t_steady']
    tau = param_dict["time_constant_exc"]
    dt = param_dict["time_step"]
    ff_scale = param_dict["feedforward_scale"]
    D = param_dict["Dimension"]
    rho = param_dict['num_neurons'] / (param_dict['position_max'] - param_dict['position_min'])
    a = param_dict['gaussian_width_exc']
    
    Rf1,Rf2 = Rf_both
    print(Rf_both)
    invCovLH = (np.sqrt(2 * np.pi) * rho) / a *np.eye(2)
    invCovLH[0,0] = invCovLH[0,0] * Rf1
    invCovLH[1,1] = invCovLH[1,1] * Rf2
    mu = param_dict['input_position']
    if isinstance(mu,int):
        mu = np.array([mu, -mu])
    muLH = mu
    prior_precision_structure = 2 * np.eye(D) - np.ones((D, D))
    invCovPost = Lambda_s * prior_precision_structure + invCovLH
    muPost = np.linalg.solve(invCovPost, invCovLH @ muLH)
    print("invCovPost", invCovPost)
    print("muPost", muPost)
    
    
    if Diag:
        Fdiag = np.diag(np.diag(invCovPost))
        rate =np.linalg.inv(Fdiag) /tau  #diagonal fisher
    else:
        Ffull = invCovPost
        Finv = np.linalg.inv(Ffull)
        print("Finv", Finv)
        rate = Finv/tau #full fisher
    print(rate)
    print("noise",sqrtm(2 * rate * dt))
    print("rate* precision", rate @ invCovPost)
    # Partially applied function for parallel execution of trials
    args = [
        (T, dt, rate, invCovPost, muPost, initial_state)
        for initial_state in initial_states_list
    ]

    with mp.get_context('spawn').Pool() as pool:
        desc = "Natural Gradient Langevin sampling with diagonal fisher" if Diag else "Natural Gradient Langevin sampling with full fisher"
        results = list(
            tqdm(
                pool.starmap(Lan_run_trial2D, args),
                total=num_trials,
                desc=desc
            )
        )
    
    return np.array(results)

def find_prior(params,num_trials, Rf_both=[10,20]):
    simulator = CANNSimulator2D(params)  # Changed to 2D simulator

    simulator.initialize_network()
    l_list = []
    kldlist = []
    for i in range(num_trials):
        Lambda_s_opt, KLD = simulator.find_prior_precision(Wee=0, Rf_both=Rf_both,normal_input=True)
        l_list.append(Lambda_s_opt)
        kldlist.append(KLD)
    print("Lambda_s_opt", l_list)
    print("KLD", kldlist)
    return(l_list[kldlist.index(min(kldlist))]),min(kldlist)
```
