## Supplementary for "Natural gradient Bayesian sampling automatically emerges in canonical cortical circuits": cannND.py

```
import numpy as np
import numpy.matlib
import matplotlib.pyplot as plt
from scipy.stats import multivariate_normal
from scipy.optimize import minimize_scalar
from matplotlib.collections import LineCollection
from tqdm import tqdm
import pickle
import multiprocessing as mp
from functools import partial
from scipy.optimize import minimize
class CANNSimulator2D:
    def __init__(self, params=None):
        """
        Initialize the CANNSimulator with given parameters.

        Parameters:
        params (dict, optional): Dictionary of parameters to override default values.
        
    A simulator for Continuous Attractor Neural Network (CANN) models.

    This class provides methods to initialize network parameters, run simulations,
    and analyze the results of CANN models. It supports various configurations
    and allows for the calibration of feedforward weights.

    Attributes:
        params (dict): Dictionary of simulation parameters.
        conn_params (dict): Dictionary of connection parameters.
        rng (np.random.Generator): Random number generator for reproducibility.

    Methods:
        initialize_network(): Initialize network connectivity.
        run_simulation(Rf, Wei, ff_scale, Wee=None, seed=None): Run the main simulation.
        calibrate_ff_scale(Rf_range=(0.1, 2), num_points=20): Calibrate feedforward weight.
        plot_precision_vs_Rf(Rf_range=(0.1, 2), num_points=2): Plot precision vs input intensity.
        analyze_results(results, Rf): Analyze simulation results.
        plot_results(results, analysis): Plot simulation results.
        compute_bump_positions_height_over_trials(Rf, Wei, ff_scale, num_trials=50, Wee=None): Compute bump positions and height over multiple trials.
        run_sim_vs_Wee_Rf(Wei, ff_scale, num_trials=50, Wee_list=None, Rf_list=None): Run simulations for different Wee and Rf values.
    """
    def __init__(self, params=None):

        default_params = {
            'time_constant_exc': 1.0,
            'position_max': 180.0,
            'position_min': -180.0,
            'gaussian_width_exc': 40.0,
            'gaussian_width_ES': 20.0,
            'num_neurons': 180,
            'simulation_time': 100.0,
            'time_step': 0.01,
            'recording_start': 20,
            'Fano_factor': 0.5,
            'normalization_k': 0.0005,
            'inhibitory_gain': 10,
            'input_position': [0,0],
            'feedforward_scale': 0.58214787,
            't_steady':20,
            'initial_mean_eq':0,
            'initial_var_eq':60, 
            'Dimension': 2
        }
        self.params = {**default_params, **(params or {})}
        self._compute_parameters()
        self.conn_params = None

        
    def _compute_parameters(self):
        """Initialize and compute all simulation parameters"""
        
        p = self.params
        #
        p['time_constant_som'] = p['time_constant_exc'] * 5

        p['gaussian_width_SE'] = np.sqrt(p['gaussian_width_exc'] ** 2 - p['gaussian_width_ES'] ** 2)

        p['gaussian_width_SS'] = np.sqrt( (p['gaussian_width_exc'] ** 2 + p['gaussian_width_ES'] ** 2) / 2)

         # Position range = max - min
        p['position_range'] = p['position_max'] - p['position_min']
        
        # Neuron density = number / range
        p['neuron_density'] = p['num_neurons'] / p['position_range']
        
        #
          # Generate preferred stimulus points for neurons
        p['PrefStim'] = np.linspace(
            p['position_min'], p['position_max'], p['num_neurons'] + 1
        )[1:p['num_neurons'] + 1]
        
        # Maximum inhibition strength
        p['max_inhibition'] = 1 / (12 * p['neuron_density'])
        
        # Critical weight calculation
        numerator = 8 * np.sqrt(2 * np.pi) * p['normalization_k'] * p['gaussian_width_exc']
        p['critical_weight'] = np.sqrt(numerator / p['neuron_density'])
        wie = 0.5 * p['critical_weight']
        p['E to S'] = np.array([[wie,wie]])
      
        # Recurrent connection weights
        p['recurrent_weight_e2e'] = 0.5 * p['critical_weight']  # E->E
        p['recurrent_weight_e2i'] = 0.5 * p['critical_weight']  # E->I
        
        # Feedforward intensity calculation
        denominator = 2 * np.sqrt(np.pi) * p['normalization_k'] * p['gaussian_width_exc']
        p['feedforward_intensity_scale'] =  (p['critical_weight'] / denominator) #Ufd
        

    def initialize_network(self):
        """Initialize network connectivity"""

        param_dict = self.params
        PrefStim = param_dict["PrefStim"]
        a_SE = param_dict["gaussian_width_SE"]
        xrange = param_dict['position_range']
        
        aE = param_dict["gaussian_width_exc"]

        a_ES = param_dict['gaussian_width_ES']


        ## Connection kernels
        W_angle = np.angle(np.exp((1j*(PrefStim-PrefStim[0])*(2*np.pi/xrange)))) * xrange/(2*np.pi)
        W_kerFtE = np.exp(-W_angle**2/(2*aE**2))/(np.sqrt(2*np.pi)*aE)    
        WE = np.expand_dims(W_kerFtE,axis=1)
        WE = np.fft.fft(W_kerFtE)
        
        W_kerFtES = np.exp(-W_angle**2/(2*a_ES**2))/(np.sqrt(2*np.pi)*a_ES)   
        WES = np.expand_dims(W_kerFtES,axis=1) 
        WES = np.fft.fft(W_kerFtES)
        
        W_kerFtSE = np.exp(-W_angle**2/(2*a_SE**2))/(np.sqrt(2*np.pi)*a_SE) 
        WSE = np.expand_dims(W_kerFtSE,axis=1)   
        WSE = np.fft.fft(W_kerFtSE)
        
        # figc1, axc1 = plt.subplots(figsize=(8,8))
        # figc1.subplots_adjust(left=0.15, bottom=0.2, right=0.85, top=0.9, wspace=0.04, hspace=0.3)
        # axc1.plot(PrefStim,W_kerFtE, label="Recurrent Kernel",linewidth=1)
        # axc1.plot(PrefStim,W_kerFtES, label="WES Kernel",linewidth=1)
        # axc1.plot(PrefStim,W_kerFtSE, label="WSE Kernel",linewidth=1)
        # axc1.legend()

        
        self.conn_params =  {"E Kernel": WE,'ES Kernel':WES, 'SE Kernel':WSE}
    def _temporal_ff(self, Rf1,Rf2,T,mu):
        """Calculate temporal feedforward input"""
        param_dict = self.params
        a = self.params["gaussian_width_exc"]
        xrange = self.params['position_range']
        PrefStim = self.params['PrefStim']
        # mu = self.params['input_position']
        # T = self.params['simulation_time']
        dt = self.params['time_step']
        N = self.params['num_neurons']
        if isinstance(mu, int):
            mu = np.array([mu, -mu])
        mu1,mu2 = mu

        x1 =  np.repeat(mu1,N)
        x2 = np.repeat(mu2,N)
        
        
        pos1 = np.subtract(x1,PrefStim)
        pos1 = np.angle(np.exp(1j*pos1 * (2*np.pi)/xrange)) * xrange/(2*np.pi)
        pos1 = np.expand_dims(pos1,axis=1)
        pos1 = np.matlib.repmat(pos1,1,int(T/dt))
        Ipos1 = Rf1 * np.exp(-(pos1**2) / (4*a**2))
        
        pos2 = np.subtract(x2,PrefStim)
        pos2 = np.angle(np.exp(1j*pos2 * (2*np.pi)/xrange)) * xrange/(2*np.pi)
        pos2 = np.expand_dims(pos2,axis=1)
        pos2 = np.matlib.repmat(pos2,1,int(T/dt))
        Ipos2 = Rf2 * np.exp(-(pos2**2) / (4*a**2))
        
        
        I1 = np.expand_dims(Ipos1,axis=1)
        I1 = np.insert(I1,1,Ipos2,axis=1)
        return I1   

    
    def run_simulation(self, Rf_both, Wei, ff_scale, Wcoup, Wee,normal_input,T = 100, noise = True):
        """Main simulation function"""
        param_dict = self.params
        
        # wse = param_dict['E to S']
        D = param_dict['Dimension']

        N = param_dict["num_neurons"]
       # T = param_dict["simulation_time"]
        dt = param_dict["time_step"]
        a = param_dict["gaussian_width_exc"]
        k = param_dict["normalization_k"]
        gain = param_dict["inhibitory_gain"]
        rho = param_dict["neuron_density"]
        F = param_dict["Fano_factor"]
        mu = param_dict['input_position']
        t_steady = param_dict['t_steady']
        tauE = param_dict["time_constant_exc"]
        tauS = param_dict['time_constant_som']
        wse = param_dict['E to S']

        WE = self.conn_params['E Kernel']
        WES = self.conn_params['ES Kernel']
        WSE = self.conn_params['SE Kernel']
        Wie = param_dict["recurrent_weight_e2i"]
        
        wes = np.array([[Wei, Wei]])
        ff = np.array([[ff_scale,ff_scale]])
        Wpeak = np.array([[Wee, Wcoup], [Wcoup, Wee]])
    

        WE = self.conn_params['E Kernel']
        WES = self.conn_params['ES Kernel']
        WSE = self.conn_params['SE Kernel']


        
        t = np.arange(0, T, dt)
        U = np.zeros((N, D, len(t)))
        V = np.zeros((N, D, len(t)))
        Oc = np.zeros((N, D, len(t)))
        Vc = np.zeros((N, D, len(t)))
        Rf1, Rf2 = Rf_both
        if normal_input:        
 
            Iext =self._temporal_ff(Rf1,Rf2,T,mu)
            # print(Iext[:,1])
            # print(Iext[1,:])
        else:
            initial_mean = param_dict['initial_mean_eq']
            initial_var = param_dict['initial_var_eq']
            shift = np.random.normal(initial_mean,initial_var,2) #60
            Iext1 = self._temporal_ff(Rf1,Rf2,t_steady,[int(shift[0]/2)*2,int(shift[1]/2)*2])
            Iext2 = self._temporal_ff(Rf1,Rf2,T-t_steady,mu)
            Iext = np.concatenate((Iext1,Iext2),axis=2)
        if noise == False:
            F = 0
            print("noise off")
            print(param_dict['input_position'])
        
        for i in range(len(t) - 1):
            noisevr = U[:, :, i].clip(min=0)
            
            O = Oc[:, :, i]
            Oin = Vc[:, :, i]
            Oft = np.fft.fft(O, axis=0)
            Oinft = np.fft.fft(Oin, axis=0)
        
            Fre = np.multiply(WE.reshape(N, 1), Oft)
            FRE =  np.fft.ifft(Fre, axis=0) @ Wpeak
            Frse = np.multiply(WSE.reshape(N, 1), Oft)
            FRSE = wse * np.fft.ifft(Frse, axis=0) 
            Fri = np.multiply(WES.reshape(N, 1), Oinft)
            FRI = wes * np.fft.ifft(Fri, axis=0)

            U[:, :, i + 1] = ((1 - dt / tauE) * U[:, :, i]) + (FRE * dt / tauE)  + (ff * Iext[:, :, i] * dt / tauE) + (np.sqrt(F * noisevr * dt) * np.random.randn(N, D)) #+ (FRI * dt / tauE)
            V[:, :, i + 1] = ((1 - dt / tauS) * V[:, :, i]) #+ (FRSE * (dt / tauS))
            
            Ocn = U[:, :, i + 1].clip(min=0) ** 2

            Ocd = np.multiply(k, np.sum(Ocn, axis=0))
            Oc[:, :, i + 1] = np.divide(Ocn, (Ocd + 1))
            
            Vc[:, :, i + 1] = gain * V[:, :, i]

        Oc = Oc.clip(min=0)
        
        
        return {'E_stim1': self._calculate_stimulus(Oc[:, 0, :]), 
                'I_stim1': self._calculate_stimulus(Vc[:, 0, :]), 
                'E_stim2': self._calculate_stimulus(Oc[:, 1, :]), 
                'I_stim2': self._calculate_stimulus(Vc[:, 1, :]),
                'E_bump_height1': np.sum(U[:, 0, :], axis=0) / (np.sqrt(np.pi) * 2* a * rho),
                'E_bump_height2': np.sum(U[:, 1, :], axis=0) / (np.sqrt(np.pi) * 2* a * rho),            
            'E_FR': Oc, 'I_FR': Vc, 'Synaptic_Input': U, 'I_Synaptic_Input': V,'params': self.params}

    def _calculate_stimulus(self, activity):
        """Calculate stimulus position from activity"""
        pref_factor = self.params['PrefStim'] * np.pi / 180
        e = np.exp(1j * pref_factor) @ activity
        return np.angle(e, deg=True)
    
    def find_prior_precision(self,Wei = 0, Wee = 0, Rf_both = [10, 20], normal_input=False, Wcoup = None):
        """
        Find the prior precision stored in the neural network.

        This method finds Lambda_s such that the KL divergence between the
        computed posterior distribution (given by network dynamics and likelihood)
        and the sample distribution is minimized.

        Parameters:
            Wee (float): Recurrent weight for excitatory-to-excitatory connections.
            Rf_both (list, optional): List containing feedforward input intensities for two populations. Default is [10, 20].
            normal_input (bool, optional): Flag to indicate if the equilibrium test is to be performed. Default is False.

        Returns:
            Lambda_s (float): The optimized prior precision.
            KLD (float): The KL divergence between the computed posterior and the sample distribution.
        """
        # Compute the initial inverse covariance from the sample distribution.
        rho = self.params['neuron_density']
        a = self.params["gaussian_width_exc"]
        Rf1,Rf2 = Rf_both
        invCovLH = (np.sqrt(2 * np.pi) * rho) / a *np.eye(2)
        invCovLH[0,0] = invCovLH[0,0] * Rf1
        invCovLH[1,1] = invCovLH[1,1] * Rf2
        mu = self.params['input_position']
        muLH = mu#np.array([mu, -mu])
        if Wcoup is None:
            Wcoup = 0.8 * self.params['critical_weight']
        # Run the simulation to get the sample distribution for lan
        results = self.compute_bump_positions_height_over_trials(Rf_both, Wei = Wei , ff_scale = self.params['feedforward_scale'], Wcoup = Wcoup, Wee = Wee, num_trials=100, normal_input=True)
        S1 = results['bump_positions1'][:,int((self.params['recording_start']*2+self.params['t_steady'])/self.params['time_step']):].reshape(-1)
        S2 = results['bump_positions2'][:,int((self.params['recording_start']*2+self.params['t_steady'])/self.params['time_step']):].reshape(-1)
        muS = np.array([np.mean(S1), np.mean(S2)])
        covS = np.cov(np.vstack((S1, S2)))
        print("muS", muS)
        print("covS", covS)


        invCovPost0 = np.linalg.inv(covS)
        print("invCovLH", invCovLH)
        print("invCovPost0", invCovPost0)
        maxLambda_s = 2 * invCovPost0[0, 0]  
        Lambda_s0 = -invCovPost0[0, 1]   

        # Optimize Lambda_s in the range [0, maxLambda_s] using a lambda to pass the extra parameters.

        # res = minimize_scalar(lambda L: self._compute_KLD(L, muS, covS, muLH, invCovLH),
        #                     bounds=(0, maxLambda_s),
        #                     method='bounded',
        #                     options={'xtol': 1e-4})
        res = minimize(lambda L: self._compute_KLD(L, muS, covS, muLH, invCovLH),
                        x0=Lambda_s0,
                        bounds=[(0, maxLambda_s)],
                        tol=1e-6,
                        method='SLSQP',
                        options={'disp': False}
                    )
        Lambda_s_opt = res.x[0]
        KLD = self._compute_KLD(Lambda_s_opt, muS, covS, muLH, invCovLH)
        return Lambda_s_opt, KLD

    def _get_KL_div(self, mu2, cov2, mu1, cov1):
        """
        Compute the KL divergence between two multivariate Gaussian distributions.
        
        Computes KL(Q||P) where:
            Q ~ N(mu1, cov1)
            P ~ N(mu2, cov2)
        
        Parameters:
            mu1, mu2 (np.ndarray): Mean vectors.
            cov1, cov2 (np.ndarray): Covariance matrices.
        
        Returns:
            float: The KL divergence.
        """
        d = len(mu1)
        inv_cov2 = np.linalg.inv(cov2)
        diff = mu2 - mu1
        term1 = np.trace(inv_cov2 @ cov1)
        term2 = diff.T @ inv_cov2 @ diff
        term3 = np.log(np.linalg.det(cov2) / np.linalg.det(cov1))
        return 0.5 * (term1 + term2 - d + term3)

    def _compute_KLD(self, Lambda_s, muS, covS, muLH, invCovLH):
        """
        Compute the KL divergence for a given prior precision Lambda_s.

        Parameters:
            Lambda_s (float): The prior precision value.
            muS (np.ndarray): Mean of the sample (target) distribution.
            covS (np.ndarray): Covariance of the sample (target) distribution.
            muLH (np.ndarray): Mean of the likelihood distribution.
            invCovLH (np.ndarray): Precision matrix (inverse covariance) of the likelihood.

        Returns:
            float: The KL divergence between the computed posterior and the sample distribution.
        """
        n = invCovLH.shape[0]
        # Build the prior precision structure: 2*I - 1
        prior_precision_structure = 2 * np.eye(n) - np.ones((n, n))
        invCovPost = Lambda_s * prior_precision_structure + invCovLH
        # Solve for posterior mean using a linear solver for numerical stability.
        muPost = np.linalg.solve(invCovPost, invCovLH @ muLH)
        # The covariance of the posterior is the inverse of the precision matrix.
        covPost = np.linalg.inv(invCovPost)
        return self._get_KL_div(muPost, covPost, muS, covS)


    def compute_bump_positions_height_over_trials(self, Rf_both, Wei, ff_scale, Wcoup, Wee, num_trials=50, normal_input=False):
        """Computes excitatory stimulation for input conditions over multiple trials."""

        args = [
            (Rf_both, Wei, ff_scale, Wcoup, Wee, normal_input, self.params['simulation_time'],True)  # Updated args to match run_simulation params
            for _ in range(num_trials)
        ]

        with mp.get_context('spawn').Pool() as pool:
            results = list(
                tqdm(
                    pool.starmap(self.run_simulation, args),
                    total=num_trials,
                    desc=f"Running trials"
                )
            )

        all_trials_pos1 = [result['E_stim1'] for result in results]
        all_trials_pos2 = [result['E_stim2'] for result in results]
        all_trials_height1 = [result['E_bump_height1'] for result in results]
        all_trials_height2 = [result['E_bump_height2'] for result in results]

        return {
            'time': self.params['simulation_time'],
            'bump_positions1': np.array(all_trials_pos1),
            'bump_positions2': np.array(all_trials_pos2), 
            'bump_height1': np.array(all_trials_height1),
            'bump_height2': np.array(all_trials_height2)
        }
    
    def get_vector_field(self, Rf_both, Wei, ff_scale, Wcoup, Wee, xrange=np.linspace(-8, 10, 20), yrange=np.linspace(-8, 10, 20), num_steps=1):
        """
        Calculate the vector field for a neural network model based on the given parameters.

        Args:
        - Rf_both: tuple, receptive fields for both dimensions
        - Wei: float, inhibitory weight
        - ff_scale: float, feedforward scaling factor
        - Wcoup: float, coupling weight
        - Wee: float, excitatory weight
        - xrange: numpy array, range of x values
        - yrange: numpy array, range of y values
        - num_steps: int, number of time steps

        Returns:
        - v1_list: numpy array, vector field for dimension 1
        - v2_list: numpy array, vector field for dimension 2
        - Lambda_s: float, prior precision
        """
        
        result = self.run_simulation(Rf_both, Wei, ff_scale, Wcoup, Wee, normal_input=True,T = 40,noise = False)
        E_stim1 = result['E_stim1'][-1]
        E_stim2 = result['E_stim2'][-1]
        E_bump_height1 = result['E_bump_height1']
        E_bump_height2 = result['E_bump_height2']
        Oc = result['E_FR']
        Vc = result['I_FR']
        U = result['Synaptic_Input']
        V = result['I_Synaptic_Input']
        param_dict = self.params
        
        # wse = param_dict['E to S']
        D = param_dict['Dimension']

        N = param_dict["num_neurons"]
       # T = param_dict["simulation_time"]
        dt = param_dict["time_step"]
        a = param_dict["gaussian_width_exc"]
        k = param_dict["normalization_k"]
        gain = param_dict["inhibitory_gain"]
        rho = param_dict["neuron_density"]
        F = param_dict["Fano_factor"]
        mu = param_dict['input_position']
        t_steady = param_dict['t_steady']
        tauE = param_dict["time_constant_exc"]
        tauS = param_dict['time_constant_som']
        wse = param_dict['E to S']

        WE = self.conn_params['E Kernel']
        WES = self.conn_params['ES Kernel']
        WSE = self.conn_params['SE Kernel']
        Wie = param_dict["recurrent_weight_e2i"]
        
        wes = np.array([[Wei, Wei]])
        ff = np.array([[ff_scale,ff_scale]])
        Wpeak = np.array([[Wee, Wcoup], [Wcoup, Wee]])
        print("Wpeak",Wpeak)
    

        WE = self.conn_params['E Kernel']
        WES = self.conn_params['ES Kernel']
        WSE = self.conn_params['SE Kernel']


        Rf1, Rf2 = Rf_both
        mu1,mu2 = mu
        l1 = len(xrange)
        l2 = len(yrange)
        v1_list = np.zeros((l1,l2,num_steps))
        v2_list = np.zeros((l1,l2,num_steps))
        print("U",U[:, :, -1].shape)
        print("Oc",Oc[:, :, -1].shape)
        
        
        U_hist = np.zeros((N, D, num_steps + 1))
        V_hist = np.zeros((N, D, num_steps + 1))
        Oc_hist = np.zeros((N, D, num_steps + 1))
        # Initialize with the current state
        U_hist[:, :, 0] = U[:, :, -1]
        V_hist[:, :, 0] = V[:, :, -1]
        Oc_hist[:, :, 0] = Oc[:, :, -1]
        Lambda_s,kld = self.find_prior_precision(Wei = Wei, Wee=Wee, Rf_both=Rf_both,normal_input=True)
        
        print("current pos",E_stim1,E_stim2)
        Rf1,Rf2 = Rf_both
        invCovLH = (np.sqrt(2 * np.pi) * rho) / a *np.eye(2)
        invCovLH[0,0] = invCovLH[0,0] * Rf1
        invCovLH[1,1] = invCovLH[1,1] * Rf2
        mu = param_dict['input_position']
        if isinstance(mu,int):
            mu = np.array([mu, -mu])
        muLH = mu
        prior_precision_structure = 2 * np.eye(D) - np.ones((D, D))
        invCovPost = Lambda_s * prior_precision_structure + invCovLH
        coef_pre = np.linalg.inv(invCovLH)@invCovPost
        for i, n1 in enumerate(xrange):
            for j, n2 in enumerate(yrange):
                
                Iext = self._temporal_ff(Rf1, Rf2, (num_steps)* dt, np.array([mu1,mu2])+ coef_pre@ np.array([E_stim1-n1, E_stim2-n2]))
                for t in range(num_steps):


                    O = Oc_hist[:, :, t]
                    Oin = V_hist[:, :, t]

                    Oft = np.fft.fft(O, axis=0)
                    Oinft = np.fft.fft(Oin, axis=0)

                    Fre = np.multiply(WE.reshape(N, 1), Oft)
                    FRE = np.fft.ifft(Fre, axis=0) @ Wpeak
                    Frse = np.multiply(WSE.reshape(N, 1), Oft)
                    FRSE = np.fft.ifft(Frse, axis=0) * wse
                    Fri = np.multiply(WES.reshape(N, 1), Oinft)
                    FRI = np.fft.ifft(Fri, axis=0) * wes

                    newU = ((1 - dt / tauE) * U_hist[:, :, t]) + (FRE * dt / tauE) + (ff * Iext[:, :, t] * dt / tauE)


                    Ocn = newU.clip(min=0) ** 2
                    Ocd = np.multiply(k, np.sum(Ocn, axis=0))
                    newOc = np.divide(Ocn, (Ocd + 1))
                    newOc = newOc.clip(min=0)

                    U_hist[:, :, t + 1] = newU
                    Oc_hist[:, :, t + 1] = newOc

                    # Store the final values for analysis after full time evolution
                    finalOc = Oc_hist[:, :, -1]
                    v1_list[i, j,t] = self._calculate_stimulus(Oc_hist[:, 0, t + 1]) - self._calculate_stimulus(Oc_hist[:, 0, t ])
                    v2_list[i, j,t] = self._calculate_stimulus(Oc_hist[:, 1, t + 1]) - self._calculate_stimulus(Oc_hist[:, 1, t ])


        
        return v1_list, v2_list, Lambda_s

def save_simulation_data_pickle(data, filename):
    with open(filename, 'wb') as f:
        pickle.dump(data, f)   
def load_simulation_data_pickle(filename):
    with open(filename, 'rb') as f:
        return pickle.load(f)
```
