## Supplementary for "Natural gradient Bayesian sampling automatically emerges in canonical cortical circuits": plotdataND.py

```
import matplotlib.pyplot as plt
import seaborn as sns
import numpy as np
from scipy.stats import norm
from matplotlib.collections import LineCollection
from matplotlib.gridspec import GridSpec
from mpl_toolkits.axes_grid1 import make_axes_locatable
import os
import sys
import gc
from cannND import *
from analysis_tools2D import *
import os
def save_fig(fig, filename, sampling = "Langevin"):
    """
    Save the figure with a white background.
    Args:
        fig: The figure to save.
        filename: The name of the file to save the figure as.
    """
    fig.patch.set_facecolor('none')   # Set the patch (outer part) to transparent
    fig.set_facecolor('none')   # Set the facecolor of the figure to white
    # Turn off axis for all subplots
    for ax in fig.axes:
        #ax.set_axis_off()
        #ax.set_xlim(left=0)
        ax.tick_params(
        axis='both',        # Apply to both x and y
        which='both',       # Apply to both major and minor ticks
        direction='in',     # Tick direction: inward
        length=4,
        width=1,
        labelsize=10
        )
        xlabel = ax.get_xlabel()  
        ylabel = ax.get_ylabel()  

        ax.set_xlabel(xlabel, fontsize=16) 
        ax.set_ylabel(ylabel, fontsize=16)

        # Optional: make ticks thinner and closer if desired
        ax.tick_params(width=1, length=4, labelsize=10)

    if sampling == 'Hamiltonian':
        folder_path = 'Ham'
    else:
        folder_path = 'Lan'
    if not os.path.exists(folder_path):
        os.makedirs(folder_path)
    plt.rcParams['ps.fonttype'] = 42
    plt.rcParams['pdf.fonttype'] = 42
    filename = os.path.join(folder_path, filename)
    fig.savefig(filename, format='eps',transparent = True,  bbox_inches='tight')
    plt.close(fig)


def plot_one_simulationND(params, Rf_both=[10,20],normal_input=True,tstart = 0,tend = 100):
    """
    Plot one simulation for Langevin sampling

    Args:
    - params: dictionary containing simulation parameters
    - Rf_both: list, input intensity for 2D Langevin sampling
    - normal_input: bool, whether to use normal input
    - tstart: int, start time for plotting
    - tend: int, end time for plotting
    """
    simulator = CANNSimulator2D(params)
    simulator.initialize_network()
    results_Lan = simulator.run_simulation(
        Rf_both, 
        Wei = 0, 
        ff_scale=params['feedforward_scale'],
        Wcoup = 0.8*simulator.params['critical_weight'] , 
        Wee = simulator.params['recurrent_weight_e2e'],
        T = params['simulation_time'],
        normal_input= normal_input
    )
    plot_results(results_Lan, "Langevin",tstart = tstart,tend = tend)


def plot_results(results,sampling,tstart = 0,tend = 100):
    """Visualization methods"""
    plot_firing_rates(results,sampling,tstart = tstart,tend = tend)    
    plot_bump_position(results, sampling,tstart = tstart,tend = tend)
    # plot_bump_height(results,sampling,tstart = tstart,tend = tend)

def plot_firing_rates(results, sampling,tstart = 0,tend = 100):
    """Plot firing rates"""
    fig1, ax1 =  plt.subplots(figsize=(2.5,1.75))
    E_FR = results['E_FR'][:,:,int(tstart/results['params']['time_step']):int(tend/results['params']['time_step'])]
    # First row: Full time range
    im1 = ax1.imshow(E_FR[:, 0, :], aspect='auto', cmap='Blues',
        extent=[0, tend-tstart, -180, 180], origin='lower',vmin=0,vmax=100)
    ax1.set_yticks([-180, -120,-60,0,60,120,180])
    ax1.set_xlim(left=0)
    plt.colorbar(im1, ax=ax1, label='Firing Rate', shrink=0.5, ticks=[0, 50, 100])
    ax1.set_title('Population 1 - Full Time Range')
    ax1.set_xlabel('Time Steps')
    ax1.set_ylabel('Neuron Index')
    save_fig(fig1, f'Firing_Rates_2D_p1{sampling}.eps',sampling=sampling)
    fig2, ax2 =  plt.subplots(figsize=(2.5,1.75))
    im2 = ax2.imshow(E_FR[:, 1, :], aspect='auto', cmap='Blues',
        extent=[0, tend-tstart, -180, 180], origin='lower',vmin=0,vmax=100)
    ax2.set_yticks([-180, -120,-60,0,60,120,180])
    ax2.set_yticks([-180, -120,-60,0,60,120,180])
    ax2.set_xlim(left=0)
    #ax2.set_ylim(-180, 180)
    plt.colorbar(im2, ax=ax2, label='Firing Rate', shrink=0.5, ticks=[0, 50, 100])
    ax2.set_title('Population 2 - Full Time Range')
    ax2.set_xlabel('Time Steps')
    ax2.set_ylabel('Neuron Index')

    # # Second row: Steady state period only
    # t_steady_idx = int(self.params['t_steady']/self.params['time_step'])
    # im3 = ax3.imshow(E_FR[:, 0, :t_steady_idx], aspect='auto', cmap=plt.get_cmap('Blues'),
    #     extent=[0, self.params['t_steady'], 180, -180])
    # plt.colorbar(im3, ax=ax3, label='Firing Rate', shrink=0.5)
    # ax3.set_title('Population 1 - Steady State Period')
    # ax3.set_xlabel('Time Steps')
    # ax3.set_ylabel('Neuron Index')

    # im4 = ax4.imshow(E_FR[:, 1, :t_steady_idx], aspect='auto', cmap=plt.get_cmap('Blues'),
    #     extent=[0, self.params['t_steady'], 180, -180])
    # plt.colorbar(im4, ax=ax4, label='Firing Rate', shrink=0.5)
    # ax4.set_title('Population 2 - Steady State Period')
    # ax4.set_xlabel('Time Steps')
    # ax4.set_ylabel('Neuron Index')

    save_fig(fig2, f'Firing_Rates_2D_p2_{sampling}.eps',sampling=sampling)
        

def plot_bump_position(results, sampling,tstart = 0,tend = 100):
    """Plot bump position over time"""
    t = np.arange(0, tend-tstart, results['params']['time_step'])
    figbp, axbp = plt.subplots(figsize=(2.5,1.75))

    # Plot for population 1
    axbp.plot(t, results['E_stim1'][int(tstart/results['params']['time_step']):int(tend/results['params']['time_step'])])
    axbp.set_xlim(0,tend - tstart)
    axbp.set_yticks([-6, -4,-2,0,2,4,6])
    axbp.set_xticks([0,(tend-tstart)/2,tend-tstart])
    axbp.set_xlabel(r'Time /$\tau$')
    axbp.set_ylabel(r'Bump Position')
    axbp.set_title("Bump Position Over Time")
    save_fig(figbp, f'one_simulation_BP_p1_{sampling}.eps', sampling)
    plt.close()

    # Plot for population 2 
    figbp, axbp = plt.subplots(figsize=(2.5,1.75))
    axbp.set_ylim(-6,6)
    axbp.plot(t, results['E_stim2'][int(tstart/results['params']['time_step']):int(tend/results['params']['time_step'])])
    
    axbp.set_xlim(0,tend - tstart)
    axbp.set_yticks([-6, -4,-2,0,2,4,6])
    axbp.set_xticks([0,(tend-tstart)/2,tend-tstart])
    axbp.set_xlabel(r'Time /$\tau$')
    axbp.set_ylabel(r'Bump Position')
    axbp.set_title("Bump Position Over Time")
    save_fig(figbp, f'one_simulation_BP_p2_{sampling}.eps', sampling)
    plt.close()
def plot_bump_position2D(params,Rf_both,sampling,normal_input):
    """
    Plot the bump position for coupled network.

    Args:
    - params: dictionary containing simulation parameters
    - Rf_both: list, input intensity for 2D Langevin sampling
    - normal_input: bool, whether to use normal input
    - sampling (str): Sampling identifier.
    """
    in1,in2 =params['input_position']    
    simulator = CANNSimulator2D(params)
    simulator.initialize_network()
    results= simulator.run_simulation(
        Rf_both, 
        Wei = 0, 
        ff_scale=params['feedforward_scale'],
        Wcoup = 0.8*simulator.params['critical_weight'] , 
        Wee = simulator.params['recurrent_weight_e2e'],
        T = params['simulation_time'],
        normal_input= normal_input
    )
    x1 = results['E_stim1'][int(results['params']['recording_start']/results['params']['time_step']):]
    y1 = results['E_stim2'][int(results['params']['recording_start']/results['params']['time_step']):]


    data = np.vstack([x1, y1]).T

    # Calculate mean and covariance matrix
    mean = np.mean(data, axis=0)
    cov = np.cov(data.T)

    # Create grid range (slightly expanded based on data range)
    plot_range1 = 4
    plot_range2 = 6
    x = np.linspace(-plot_range1,plot_range2, 200)
    y = np.linspace(-plot_range1,plot_range2, 200)

    # Create GridSpec layout
    
    Lambda_s, kld = simulator.find_prior_precision(
        Wei = 0, 
        Wee = simulator.params['recurrent_weight_e2e'], Rf_both=Rf_both,normal_input=True)

    # posterior
    invCovPost, muPost = compute_posterior_precision(Rf_both=Rf_both, param_dict=simulator.params, Lambda_s=Lambda_s)
    rho = params['num_neurons'] / (params['position_max'] - params['position_min'])
    print("precision",invCovPost,muPost)
    # calculate 2D gaussian density for heatmap
    X, Y = np.meshgrid(np.linspace(-plot_range1, plot_range2, int(20*(plot_range2+plot_range1)*rho)), np.linspace(-plot_range1, plot_range2, int(20*(plot_range2+plot_range1)*rho)),indexing='ij')
    pos = np.dstack((X, Y))
    rv = multivariate_normal(mean=muPost, cov=np.linalg.inv(invCovPost))
    posterior = rv.pdf(pos)


    # plot

    fig = plt.figure(figsize=(6, 6))
    gs = GridSpec(4, 4)

    ax_main = fig.add_subplot(gs[1:, 0:3])
    ax_xDist = fig.add_subplot(gs[0, 0:3], sharex=ax_main)
    ax_yDist = fig.add_subplot(gs[1:, 3], sharey=ax_main)    
    # Main heatmap using imshow
    extent = [-plot_range1, plot_range2, -plot_range1, plot_range2]
    ax_main.imshow(
        posterior.T, 
        extent=extent, 
        cmap='Blues',
        aspect='auto',
        alpha=0.7
    )

    # 2) prepare your trajectory data
    # Trajectory overlay (color by time) 
    trac1 = results['E_stim1'][-int((results['params']['time_constant_exc'] * 2) / results['params']['time_step']):]
    trac2 = results['E_stim2'][-int((results['params']['time_constant_exc'] * 2) / results['params']['time_step']):]
    ct = np.linspace(0, 1, len(trac1))
    points = np.vstack([trac1, trac2]).T.reshape(-1, 1, 2)
    segments = np.concatenate([points[:-1], points[1:]], axis=1)

    lc = LineCollection(segments, cmap='plasma', norm=plt.Normalize(0, 1), linewidth=2)
    lc.set_array(ct)
    ax_main.add_collection(lc)

    ax_main.axvline(x=in1,
           color='gray',
           linestyle='--',    # two dashes
           linewidth=1)
    ax_main.axhline(y=in2,
           color='gray',
           linestyle='--',    # two dashes
           linewidth=1)
    
    
    xrange = np.linspace(-plot_range1, plot_range2, 11)
    yrange = np.linspace(-plot_range1, plot_range2, 11)
    v1list,v2list,Lambda_s = simulator.get_vector_field(Rf_both, Wei=0, ff_scale = params['feedforward_scale'], Wcoup=0.8*simulator.params['critical_weight'], Wee = simulator.params['recurrent_weight_e2e'],xrange=xrange,yrange = yrange,num_steps = 1)

    t = 0
    Uq = v1list[:,:,t]/np.max(np.max(np.max(np.abs(v1list))))
    Vq = v2list[:,:,t]/np.max(np.max(np.max(np.abs(v1list))))
    Xq,Yq  = np.meshgrid(xrange,yrange,indexing='ij')
    ax_main.quiver(
        Xq, Yq, Uq, Vq,
        angles='xy', color='teal', scale_units='xy', scale=1, width=0.003
    )
    # plt.title(f"Vector Field of CANN")
    # plt.legend()
    # plt.xlabel("x")
    # plt.ylabel("y")
    # Marginal histograms
    ax_xDist.hist(x1, bins=40, color='lightsteelblue', alpha=0.6, density=True)
    ax_yDist.hist(y1, bins=40, color='plum', alpha=0.6, density=True, orientation='horizontal')
    ax_xDist.yaxis.set_ticks_position('left')  # Optional: Ensure y-axis is on the left

    ax_xDist.spines['left'].set_visible(True)  # Show left border
        # Marginal KDE plots
    # sns.kdeplot(x=x1, ax=ax_xDist, fill=True, color='steelblue')
    # sns.kdeplot(y=y1, ax=ax_yDist, fill=True, color='purple')
    ax_xDist.plot(x, norm.pdf(x, mean[0], np.sqrt(cov[0, 0])), color='steelblue', linewidth=2)
    ax_yDist.plot(norm.pdf(y, mean[1], np.sqrt(cov[1, 1])), y, color='purple', linewidth=2)
    ax_yDist.xaxis.set_ticks_position('bottom')  # Optional: Ensure x-axis is at the bottom
    ax_yDist.spines['bottom'].set_visible(True)  # Show bottom border
    ax_xDist.set_yticks([0, 0.15, 0.3])
    ax_xDist.set_ylim(0,0.4)
    ax_yDist.set_xticks([0, 0.15, 0.3])
    ax_yDist.set_xlim(0,0.4)
    ax_xDist.set_xlim(ax_main.get_xlim())
    ax_yDist.set_ylim(ax_main.get_ylim())
    plt.setp(ax_xDist.get_xticklabels(), visible=False)
    plt.setp(ax_yDist.get_yticklabels(), visible=False)
    ax_xDist.tick_params(bottom=False)
    ax_yDist.tick_params(left=False)


    # Main plot settings
    ax_main.set_xlim(-plot_range1, plot_range2)
    ax_main.set_ylim(-plot_range1, plot_range2)
    
    fig.suptitle("Network sampling distribution", fontsize=14)
    ax_main.set_xlabel("Population 1 Position (degrees)")
    ax_main.set_ylabel("Population 2 Position (degrees)")

    cbar_ax = fig.add_axes([0.15, 0.08, 0.7, 0.03])  # [left, bottom, width, height]
    cbar = fig.colorbar(lc, cax=cbar_ax, orientation='horizontal')
    cbar.set_label('Elapsed time')
    cbar.set_ticks([0, 1])
    cbar.set_ticklabels(['0', r'2 $\tau'])
    plt.tight_layout(rect=[0, 0, 1, 0.95]) 

    save_fig(fig, f'one_simulation_2D_BP_2D_{sampling}.eps', sampling)
    plt.close()


def plot_bump_height(results, sampling):
    """Plot bump height over time"""
    t = np.arange(0, results['params']['simulation_time'], results['params']['time_step'])
    figbp, (ax1, ax2) = plt.subplots(2, 1, figsize=(8, 12))
    figbp.subplots_adjust(left=0.1, bottom=0.1, right=0.9, top=0.9, hspace=0.3)

    # Plot for population 1
    ax1.scatter(t, results['E_bump_height1'], alpha=0.2, s=2)
    ax1.set_title("Bump Height Over Time - Population 1")
    ax1.set_xlabel("Time") 
    ax1.set_ylabel("Height")

    # Plot for population 2
    ax2.scatter(t, results['E_bump_height2'], alpha=0.2, s=2)
    ax2.set_title("Bump Height Over Time - Population 2")
    ax2.set_xlabel("Time")
    ax2.set_ylabel("Height") 
    
    save_fig(figbp, f'one_simulation_2D_BH_{sampling}.eps', sampling)


def plot_kl_divergence2D(param_dict,kl_dict, cmap,name = 'kl',T = False,sampling = 'Langevin'):
    """
    Plots the Kullback-Leibler divergence and cross-correlation of neural responses.
    Args:
        kl_dict: Dictionary containing KL divergences for different methods
    """
    if T == False:
        T = param_dict["simulation_time"]
    dt = param_dict["time_step"]
    t = np.arange(0, T+dt,dt)  

    # Create a figure with two subplots side by side
    fig, ax1 = plt.subplots(figsize=(2.5,1.75))
    for i, (method, kl) in enumerate(kl_dict.items()):
        ksim = kl / kl[0]
        ax1.semilogy(t, ksim, label=method, color=cmap[i])  # Changed to semilogy
        ax1.set_ylabel('Normalized KL divergence (log scale)')
    ax1.set_title('Normalized') 
    ax1.set_xlabel(r'Time /$\tau$')
    ax1.legend()
    save_fig(fig,f'Normalized KL Divergence of {name}.eps',sampling=sampling)

    # Plot KL divergence
    fig, ax2 = plt.subplots(figsize=(2.5,1.75))
    for i, (method, kl) in enumerate(kl_dict.items()):
        ksim = kl
        ax2.semilogy(t, ksim, label=method, color=cmap[i])  # Changed to semilogy
        ax2.set_ylabel('KL divergence (log scale)')
    ax2.set_title('KL divergence')
    ax2.set_xlabel(r'Time /$\tau$')
    ax2.legend()
    save_fig(fig, f'KL Divergence of {name}.eps',sampling=sampling)

def Get_Kl_div_vs_Lan_eq2D(params,Wee = 0, Rf_both = [10,20], num_trials=50,normal_input = False,Lambda_s_opt = 0.36677033509467311):
    """
    Computes and plots the Kullback-Leibler divergence for different methods 
    and analyzes the bump positions and heights in a 2D neural network simulation.

    Args:
        params (dict): Dictionary containing simulation parameters.
        Wee (float): Recurrent weight for excitatory-to-excitatory connections.
        Rf_both (list): List containing receptive field sizes for two populations.
        num_trials (int): Number of trials for the simulation.
        normal_input (bool): Whether to test equilibrium conditions.

    Returns:
        None
    """

    simulator = CANNSimulator2D(params)  # Changed to 2D simulator
    simulator.initialize_network()
    #Lambda_s_opt, KLD = simulator.find_prior_precision(Wee=Wee, Rf_both=Rf_both,normal_input=True)
  #  print(Lambda_s_opt,KLD)
    
    kl_dict = {}
    se_dict = {}
    Ue1_list = []

    T = params['simulation_time']

    print(Rf_both)
    results = simulator.compute_bump_positions_height_over_trials(
        Rf_both, Wei=0, ff_scale=params['feedforward_scale'], Wcoup = 0.8*simulator.params['critical_weight'] , Wee=Wee, num_trials=num_trials, normal_input=False
    )
    Rf1,Rf2 = Rf_both           
    Se_Lan_CANN1 = results['bump_positions1'][:, int(params['t_steady'] / params['time_step'])-1:]
    Se_Lan_CANN2 = results['bump_positions2'][:, int(params['t_steady'] / params['time_step'])-1:]

    kl_Lan_CANN = compute_kl_divergence2D(params, Lambda_s_opt, Se_Lan_CANN1, Se_Lan_CANN2, Rf_both, num_trials=num_trials)
    kl_dict[f'CANN'] = kl_Lan_CANN
    se_dict[f'CANN'] = np.concatenate((Se_Lan_CANN1,Se_Lan_CANN2),axis = 0)    

    initials_states_list1 = results['bump_positions1'][:,int(params['t_steady']/params['time_step'])-1]
    initials_states_list2 = results['bump_positions2'][:,int(params['t_steady']/params['time_step'])-1]
    initials_states_list = np.column_stack((initials_states_list1,initials_states_list2))
    Lan_full =  langevin_sampling2D(params, Rf_both,Lambda_s_opt, initials_states_list, num_trials=num_trials, normal_input=False, Diag=False)
    kl_full = compute_kl_divergence2D(params, Lambda_s_opt, Lan_full[:,:,0], Lan_full[:,:,1], Rf_both, num_trials=num_trials)
    
    Lan_diag =  langevin_sampling2D(params, Rf_both,Lambda_s_opt, initials_states_list, num_trials=num_trials, normal_input=False, Diag=True)
    kl_diag= compute_kl_divergence2D(params, Lambda_s_opt, Lan_diag[:,:,0], Lan_diag[:,:,1], Rf_both, num_trials=num_trials)

    kl_dict[f'Natural Gradient (Diagonal)'] = kl_diag
    se_dict[f'Natural Gradient (Diagonal)'] = Lan_diag
    kl_dict[f'Natural Gradient (Full)'] = kl_full
    se_dict[f'Natural Gradient (Full)'] = Lan_full
    cmap = plt.cm.spring(np.linspace(0, 1, len(kl_dict)))
    plot_kl_divergence2D(param_dict=params, kl_dict=kl_dict, cmap =cmap, name='2D', T=params['simulation_time'] - params['t_steady'],sampling='Langevin')


def plot_vector_field_LAN(param_dict, Rf_both, Wei, ff_scale, Wee, stride=1, Lambda_s=None, fisher='both'):
    """
    Plots vector fields (diagonal and/or full Fisher) over a posterior heatmap for a 2D CANN.

    Args:
        param_dict (dict): Network parameters, must include:
            'Dimension', 'num_neurons', 'position_max', 'position_min',
            'gaussian_width_exc', and 'input_position'.
        Rf_both (tuple): (Rf1, Rf2) - scaling factors for the likelihood precision along x and y.
        Wei: Feedforward weights (unused).
        ff_scale: Feedforward scale (unused).
        Wee: Recurrent weight for prior precision.
        stride (int): Stride for subsampling the vector field grid.
        Lambda_s (float, optional): Prior precision; if None, it will be computed.
        fisher (str): 'diag', 'full', or 'both' to control which Fisher vector field(s) to plot.
    """
    # Extract key parameters
    D = param_dict["Dimension"]
    rho = param_dict['num_neurons'] / (param_dict['position_max'] - param_dict['position_min'])
    a = param_dict['gaussian_width_exc']

    # Initialize simulator and compute prior precision if needed
    
    simulator = CANNSimulator2D(param_dict)
    simulator.initialize_network()
    if Lambda_s is None:
        Lambda_s, _ = simulator.find_prior_precision(Wei=Wei, Wee=Wee, Rf_both=Rf_both, normal_input=True)

    # Grid setup
    plot_range1 = 4
    plot_range2 = 6
    xrange = np.linspace(-plot_range1, plot_range2, 11)
    yrange = np.linspace(-plot_range1, plot_range2, 11)
    Xq, Yq = np.meshgrid(xrange[::stride], yrange[::stride])

    invCovPost, muPost = compute_posterior_precision(Rf_both, param_dict, Lambda_s)

    # Posterior heatmap
    res = int(20 * (plot_range2+plot_range1) * rho)
    Xh, Yh = np.meshgrid(
        np.linspace(-plot_range1, plot_range2, res),
        np.linspace(-plot_range1, plot_range2, res),indexing='ij')
    pos = np.dstack((Xh, Yh))
    rv = multivariate_normal(mean=muPost, cov=np.linalg.inv(invCovPost))
    posterior = rv.pdf(pos)

    # Plot heatmap
    plt.figure(figsize=(4,4))
    plt.imshow(posterior, extent=[-plot_range1, plot_range2, -plot_range1, plot_range2],
               origin='lower',
                cmap='Blues',
                aspect='auto',
                alpha=0.7)
    plt.colorbar(label='Posterior Probability Density')
    mu = param_dict['input_position']
    in1, in2 = mu
    plt.axvline(x=in1,
           color='gray',
           linestyle='--',    # two dashes
           linewidth=1)
    plt.axhline(y=in2,
           color='gray',
           linestyle='--',    # two dashes
           linewidth=1)
    # Determine which Fisher types to plot
    fisher_types = ['diag', 'full'] if fisher == 'both' else [fisher]

    for fisher_type in fisher_types:
        if fisher_type == 'diag':
            fisher_inv = np.linalg.inv(np.diag(np.diag(invCovPost)))
            color = 'teal'
        elif fisher_type == 'full':
            fisher_inv = np.linalg.inv(invCovPost)
            color = 'darkorange'
        else:
            raise ValueError(f"Invalid fisher type: {fisher_type}")

        # Compute vector field
        U, V = np.zeros_like(Xq), np.zeros_like(Yq)
        for i in range(U.shape[0]):
            for j in range(U.shape[1]):
                pos_ij = np.array([xrange[i * stride], yrange[j * stride]])
                grad = (fisher_inv @ invCovPost) @ (muPost - pos_ij)
                U[i, j] = grad[0]
                V[i, j] = grad[1]

        # Plot quiver
        plt.quiver(Xq, Yq, U / 5, V / 5,
        angles='xy', scale_units='xy', scale=1,
        color=color, width=0.003, label=fisher_type)


    plt.title(f"Vector Field: {fisher} Fisher")
    plt.xlabel('x')
    plt.ylabel('y')
    plt.legend()
    plt.gca().set_aspect('equal', 'box')

    # Save the figure
    filename = f"VectorField_diag_{fisher}_{Rf_both}.eps"
    save_fig(plt.gcf(), filename)


def plot_precision_prior(params,num_trials =50):
    """
    Plot the precision prior by running simulations and comparing theoretical vs experimental means and precisions.

    Args:
    params (dict): Dictionary containing simulation parameters.
    num_trials (int): Number of simulation trials to run. Default is 50.
    """

    simulator = CANNSimulator2D(params)
    simulator.initialize_network()
    intheo = []
    pretheo =[]
    inexp =[]
    preexp = []
    for i in range(num_trials):
        Wcoup = np.random.uniform(0, simulator.params['critical_weight'])
        
        mu = [np.random.uniform(-5,5),np.random.uniform(-5,5)]

        simulator.params['input_position'] =mu
        Wee = np.random.uniform(0, simulator.params['critical_weight'])
        Rf_both = [np.random.uniform(5,20),np.random.uniform(5,20)]
        Lambda_s, kld = simulator.find_prior_precision(Wei = 0, Wee = Wee ,Rf_both=Rf_both,normal_input=True,Wcoup=Wcoup)
        results = simulator.run_simulation(
            Rf_both=Rf_both, Wei=0, ff_scale=params['feedforward_scale'],
            Wcoup=Wcoup, Wee=Wee,
            T=params['simulation_time'],
            normal_input=False
        )
        bp1 = results['E_stim1'][int(params['recording_start']/params['time_step']):]
        bp2 = results['E_stim2'][int(params['recording_start']/params['time_step']):]

        # Combine into a two-dimensional array, shape = (n_samples, 2)
        data = np.vstack([bp1, bp2]).T
        # Calculate the mean and covariance matrix
        mean = np.mean(data, axis=0)
        for j in mean:
            inexp.append(j)
        # inexp.append(mean)
        cov = np.cov(data.T)
        print(cov,np.linalg.inv(cov))
        preexp.append(np.linalg.inv(cov)[0,1])
        invCovPost, muPost = compute_posterior_precision(Rf_both=Rf_both, param_dict=simulator.params, Lambda_s=Lambda_s)
        pretheo.append(invCovPost[0,1])
        print(invCovPost)
        for j in muPost:
            intheo.append(j)
    np.save(f'precision_prior{num_trials}.npy', {
    'intheo': intheo,
    'inexp': inexp,
    'pretheo': pretheo,
    'preexp': preexp})

    plt.figure(figsize=(4, 4))
    plt.scatter(intheo, inexp)
    plt.plot(np.arange(-5,5,1), np.arange(-5,5,1), color='black',linestyle='--')
    plt.xlabel('Theoretical mean')
    plt.ylabel('Experimental mean')
    save_fig(plt.gcf(), f"Mean_prior{num_trials}.eps",sampling='Langevin')
    plt.figure(figsize=(4, 4))
    plt.scatter(-np.array(pretheo), -np.array(preexp))
    plt.plot(np.linspace(-0.1,0.8,3), np.linspace(-0.1,0.8,3), color='black',linestyle='--')
    plt.xlabel('Theoretical precision')
    plt.ylabel('Experimental precision')
    save_fig(plt.gcf(), f"Precision_prior{num_trials}.eps",sampling='Langevin')


def plot_precision_prior_fromdic(filename = 'precision_prior2.npy'):
    # Load the data from the .npy file
    # allow_pickle=True is required to load objects like dictionaries
    loaded_data = np.load(filename, allow_pickle=True)


    data_dict = loaded_data.item()

    intheo = data_dict['intheo']
    inexp = data_dict['inexp']
    pretheo = data_dict['pretheo']
    preexp = data_dict['preexp']

    # plt.figure(figsize=(4, 4))
    # plt.scatter(intheo, inexp)
    # plt.plot(np.arange(-5,5,1), np.arange(-5,5,1), color='black',linestyle='--')
    # plt.xlabel('Theoretical mean')
    # plt.ylabel('Experimental mean')
    # save_fig(plt.gcf(), f"Mean_prior2.eps",sampling='Langevin')
    plt.figure(figsize=(4, 4))
    plt.scatter(-np.array(pretheo), -np.array(preexp))
    plt.plot(np.linspace(-0.1,0.8,3), np.linspace(-0.1,0.8,3), color='black',linestyle='--')
    plt.xlabel('Theoretical precision')
    plt.ylabel('Experimental precision')
    save_fig(plt.gcf(), f"Precision_prior2.eps",sampling='Langevin')


def plot_diagonal_prior(lambda_s):
    """
    Plots the joint prior p(s, z) proportional to exp[-Lambda_s (s - z)^2 / 2]
    and its marginals p(s) and p(z).

    Args:
        lambda_s (float): The prior precision.
    """
    # Define the range for stimulus (s) and context (z)
    range_val = np.linspace(-180, 180, 200)
    s, z = np.meshgrid(range_val, range_val)


    difference = s - z
    # Calculate the joint prior
    # The joint prior is proportional to exp[-lambda_s * (s - z)^2 / 2]
    # The angular difference is calculated to ensure periodicity
    angular_difference = (difference + 180) % 360 - 180
    prior_joint = np.exp(-lambda_s * angular_difference**2 / 2)

    # Create the figure and subplots
    fig,ax_joint = plt.subplots(figsize=(6, 6))
    # gs = fig.add_gridspec(3, 3, hspace=0.3, wspace=0.3)

    # ax_joint = fig.add_subplot(gs[0:2, 0:2])
    # ax_marg_s = fig.add_subplot(gs[2, 0:2], sharex=ax_joint)
    # ax_marg_z = fig.add_subplot(gs[0:2, 2], sharey=ax_joint)

    # Plot the joint prior
    im = ax_joint.imshow(prior_joint, extent=[-180, 180, -180, 180], origin='lower', aspect='auto', cmap='Blues')
    ax_joint.set_yticks([-180, -120,-60,0,60,120,180])
    ax_joint.set_xticks([-180, -120,-60,0,60,120,180])
    ax_joint.set_xlabel("Stimulus (s)")
    ax_joint.set_ylabel("Context (z)")
    ax_joint.set_title("Joint prior p(s, z)")

    # Add a colorbar
    cbar = fig.colorbar(im, ax=[ax_joint])
    cbar.set_label("Probability")

    # Add the diagonal lines (optional, for visual reference)
    # ax_joint.plot([-180, 180], [-180, 180], 'k--')
    # ax_joint.plot([-150, 150], [-180, 120], 'k--', alpha=0.5)
    # ax_joint.plot([0, 150], [-150,180 ], 'k--', alpha=0.5)

    plt.savefig(f"Diagonal_Prior_{lambda_s}.png")
    save_fig(fig, f"Diagonal_Prior_{lambda_s}.eps",sampling='Langevin')

def plot_prior_L_vs_wcoup(params):

    simulator = CANNSimulator2D(params)
    simulator.initialize_network()
    Lambda_s_list = []
    Wcoup_list = np.linspace(0, 1, 11)*simulator.params['critical_weight']

    for Wcoup in Wcoup_list:
        Lambda_s, kld = simulator.find_prior_precision(Wei=0, Wcoup=Wcoup, Rf_both=[10, 20], normal_input=True)
        Lambda_s_list.append(Lambda_s)
    np.save(f'Lambda_s_vs_wcoup.npy', Lambda_s_list)
    np.load(f'Lambda_s_vs_wcoup.npy')
    plt.figure(figsize=(4, 4))
    plt.plot(Wcoup_list, Lambda_s_list, marker='o')
    plt.xlabel(r'$W_{coup}$')
    plt.ylabel(r'$\Lambda_s$')
    plt.title('Prior Precision vs Coupling Weight')
    save_fig(plt.gcf(), f"Prior_L_vs_wcoup.eps",sampling='Langevin')
    plt.savefig(f"Prior_L_vs_wcoup.png")
```
