## Supplementary for "Natural gradient Bayesian sampling automatically emerges in canonical cortical circuits": draft_1.py

```
import numpy as np
import torch
import matplotlib.pyplot as plt
from pathlib import Path
from scipy.stats import gaussian_kde
import matplotlib.gridspec as gridspec
from matplotlib.collections import LineCollection

def plot_ze_p_joint(
        traj: np.ndarray,
        time_step: float,
        n_show: int ,
        save_path: str,
        cmap: str = "plasma",
        bins: int = 40,
        title: str = "2-D sampling distribution"):
    
    traj_out = traj.copy()
    if n_show is not None and n_show < len(traj):
        traj = traj[-n_show:]

    # -----------------------------------------------------------
    #  Filter out NaN / inf before any plotting
    # -----------------------------------------------------------
    finite_mask = np.isfinite(traj).all(axis=1)
    
    traj = traj[finite_mask]
    if traj.shape[0] < 2:                           
        print("Warning: too few finite points to plot.")  
        return                                       

    z = traj[:, 0].astype(float)
    p = traj[:, 1].astype(float)
    z_0 = traj_out[:, 0].astype(float)
    p_0 = traj_out[:, 1].astype(float)
    print(traj_out.shape)
    t = np.arange(len(traj)) * time_step

    fig = plt.figure(figsize=(4.8, 4.2))
    gs  = gridspec.GridSpec(2, 2, width_ratios=[4, 1.2],
                                   height_ratios=[1.2, 4],
                                   hspace=0.05, wspace=0.05)

    ax_main   = fig.add_subplot(gs[1, 0])
    ax_xmarg  = fig.add_subplot(gs[0, 0], sharex=ax_main)
    ax_ymarg  = fig.add_subplot(gs[1, 1], sharey=ax_main)

# -------- 2‑D Gaussian heat‑map background --------------------
    try:
        mu_vec  = np.array([z_0.mean(), p_0.mean()])     # mean
        cov_mat = np.cov(z_0, p_0)                       # covariance

        std_z, std_p = np.sqrt(np.diag(cov_mat))
        z_grid = np.linspace(mu_vec[0] - 3*std_z, mu_vec[0] + 3*std_z, 150)
        p_grid = np.linspace(mu_vec[1] - 3*std_p, mu_vec[1] + 3*std_p, 150)
        xi, yi = np.meshgrid(z_grid, p_grid)

        from scipy.stats import multivariate_normal
        rv = multivariate_normal(mean=mu_vec, cov=cov_mat)
        zi = rv.pdf(np.dstack((xi, yi)))                 # density
        extent = [z_grid.min(), z_grid.max(), p_grid.min(), p_grid.max()]
        # 归一化到 [0,1] 方便调 alpha
        zi = (zi - zi.min()) / (zi.max() - zi.min() + 1e-9)


        contour_levels = np.linspace(0.05,1, 5)  # 可以自定义等高线等级
        ax_main.contour(
            xi, yi, zi,
            levels=contour_levels,
            cmap="Blues",
            linewidths=1.2,
            alpha=0.8
        )


    except Exception as e:
        print("Heat‑map skipped:", e)
    # -------- Trajectory coloured by time ----------------------
    points = np.array([z, p]).T.reshape(-1, 1, 2)
    segments = np.concatenate([points[:-1], points[1:]], axis=1)

    norm = plt.Normalize(t.min(), t.max())
    lc = LineCollection(segments, cmap=cmap, norm=norm,linewidth=1.0,alpha=0.6)
    lc.set_array(t[:-1])       
    lc.set_linewidth(1.0)
    lc.set_alpha(0.9)

    line = ax_main.add_collection(lc)
    cbar = fig.colorbar(line, ax=ax_main, fraction=0.046, pad=0.04)
    cbar.set_label("Elapsed time")

    # cbar_bg_ax = fig.add_axes([0.92, 0.11, 0.02, 0.70])  # [left, bottom, width, height]
    # cbar_bg = fig.colorbar(contour, cax=cbar_bg_ax)
    # cbar_bg.set_label("Normalized density", labelpad=4)


    # -------- Marginal histograms ------------------------------
#     hist_z, bins_z = np.histogram(z_0, bins=bins, range=extent[:2])
#     hist_p, bins_p = np.histogram(p_0, bins=bins, range=extent[2:])


#     max_height = max(hist_z.max(), hist_p.max())
#     hist_z = hist_z / max_height
#     hist_p = hist_p / max_height


#     bin_centers_z = 0.5 * (bins_z[:-1] + bins_z[1:])
#     bin_centers_p = 0.5 * (bins_p[:-1] + bins_p[1:])

#     ax_xmarg.bar(bin_centers_z, hist_z, width=(bins_z[1] - bins_z[0]),
#                 color="steelblue", alpha=0.6)
#     ax_ymarg.barh(bin_centers_p, hist_p, height=(bins_p[1] - bins_p[0]),
#                 color="mediumpurple", alpha=0.6)


    # # Optional 1-D KDE curves
    # try:
    #     kde_z = gaussian_kde(z_0)
    #     xx = np.linspace(*extent[:2], 400)
    #     kde_z_vals = kde_z(xx)
    #     kde_z_vals = kde_z_vals / kde_z_vals.max() * (hist_z.max() )
    #     ax_xmarg.plot(xx, kde_z_vals, c="k", lw=1)
    # except Exception:
    #     pass

    # try:
    #     kde_p = gaussian_kde(p_0)
    #     yy = np.linspace(*extent[2:], 400)
    #     kde_p_vals = kde_p(yy)
    #     kde_p_vals = kde_p_vals / kde_p_vals.max() * (hist_p.max() )
    #     ax_ymarg.plot(kde_p_vals, yy, c="k", lw=1)
    # except Exception:
    #     pass


    ax_xmarg.axis("off")
    ax_ymarg.axis("off")
        # ② 统一坐标轴
    ax_main.set_yticks([-110,0,110])
    ax_main.set_xlabel(r"Stimulus sample $z_E$")
    ax_main.set_ylabel(r"Momentum $p$")
    ax_main.set_title(title, pad=4, fontsize=10)
    ax_main.plot(np.mean(z_0), np.mean(p_0), "o", color="green", markersize=4, label="center")
    # ax_main.plot(z[0], p[0], "o", color="green", markersize=4, label="start")
    # ax_main.plot(z[-1], p[-1], "o", color="red", markersize=4, label="end")
    ax_main.legend(loc="best", fontsize=8)
    # ax_xmarg.set_xscale("function", functions=(forward, inverse))
    # ax_ymarg.set_yscale("function", functions=(forward, inverse))
    # ax_main.set_yscale("function", functions=(forward, inverse))
    # ax_main.set_xscale("function", functions=(forward, inverse))
    # ax_main.set_ylim(forward(extent[2]), forward(extent[3]))
    # ax_main.set_xlim(forward(extent[0]), forward(extent[1]))
    # ax_main.set_xlim(-0.1,0.1)
    # ax_main.set_ylim(-0.005,0.005)


    if save_path:
        Path(save_path).parent.mkdir(parents=True, exist_ok=True)
        # plt.savefig(save_path, dpi=300, bbox_inches="tight")
        plt.savefig(save_path, dpi=300, bbox_inches="tight",format='eps')

    plt.show()
    plt.close()
def m2_to_tau(M2: torch.Tensor,
              Lambda: float = 1.0,
              Lambda_p: float = 1.0):
    """
    Recover   tau_L^{-1}, tau_H^{-1}, tau_p^{-1}
    from matrix entries of  M2  under
        m11 = -tau_L^{-1} * Lambda
        m12 =  tau_H^{-1} * Lambda_p
        m21 =  tau_H^{-1} * Lambda
        m22 = -tau_p^{-1} * Lambda_p
    """
    m11, m12 = M2[0, 0].item(), M2[0, 1].item()
    m21, m22 = M2[1, 0].item(), M2[1, 1].item()

    tau_L_inv = -m11 / Lambda
    tau_H_inv =  m12 / Lambda_p          # identical result with  m21 / Lambda
    tau_p_inv = -m22 / Lambda_p

    return tau_L_inv, tau_H_inv, tau_p_inv

def main():
    # load data
    M_2 = np.load('M_2_2.0.npy')[:,:,0]
    alpha_1 = np.load('alpha_1_2.0.npy')
    alpha_2 = np.load('alpha_2_2.0.npy')
    sigma_E = np.load('Sigma_E_2.0.npy')
    print(M_2,sigma_E[0], alpha_1[0])
    
    
    T_martic = torch.tensor([[1,0],[alpha_1[0],alpha_2[0]]]).double()
    z_sample = np.load('long_trial_outputs/long_trial_Rf_11.00.npy')
    z_sample = torch.tensor(z_sample).double()
    print(z_sample.shape)
    z_sample_0 = z_sample[2000:,:]
    traj = torch.einsum('ij,kj->ki',T_martic,z_sample_0).numpy()
    plot_ze_p_joint(traj, 0.01, 800, save_path='ze_p_joint_distribution_cann_8_tau.eps')

if __name__ == "__main__":
    main()
```
