## Supplementary for "Natural gradient Bayesian sampling automatically emerges in canonical cortical circuits": height.py

```
import numpy as np
import matplotlib.pyplot as plt
import HAM_cann_simulator as HAM
import multiprocessing as mp
import json
import os
def height_ham(args):
    params, Rf = args
    simulator = HAM.CANNSimulator(params)
    ZE, ZS, bump_height = HAM.bump_position_ham(params, Rf=Rf)
    return Rf, bump_height[2000:]


if __name__ == "__main__":
    params = {
        'time_constant_exc': 1.0,
        'position_max': 180.0,
        'position_min': -180.0,
        'gaussian_width_exc': 40.0,
        'gaussian_width_ES': 20.0,
        'num_neurons': 180,
        'simulation_time': 10000.0,
        'time_step': 0.01,
        'recording_start': 20,
        'Fano_factor': 0.5,
        'normalization_k': 0.0005,
        'inhibitory_gain': 10,
        'input_position': 0,
        'feedforward_scale': 1.16429574032,
        't_steady': 20,
        'initial_mean_eq': 0,
        'initial_var_eq': 60,
        'initial_scale_eq': 1e-1
    }
    Rf_values = [r for r in range(0, 1)]
    args_list = [(params, Rf) for Rf in Rf_values]

    with mp.Pool(processes=10) as pool:
        results = pool.map(height_ham, args_list)

    # save as dict
    height_dict = {Rf: height_array for Rf, height_array in results}

    # save as numpy folder
    os.makedirs("height_outputs", exist_ok=True)
    for Rf, arr in height_dict.items():
        filename = f"height_outputs/height_Rf_{Rf:.2f}.npy"
        np.save(filename, arr)
        print(f"Saved {filename}, shape = {arr.shape}")
    
    
```
