## Supplementary for "Natural gradient Bayesian sampling automatically emerges in canonical cortical circuits": LampdaP.py

```
import re
import numpy as np
import matplotlib.pyplot as plt
import os

plt.rcParams.update({
    "font.family": "serif",
    "font.serif": ["Computer Modern"],
    "text.usetex": False,
    "axes.labelsize": 12,
    "font.size": 11,
    "legend.fontsize": 10,
    "xtick.labelsize": 10,
    "ytick.labelsize": 10,
    "lines.linewidth": 2
})

def parse_file(filename):
    with open(filename, 'r') as f:
        content = f.read()

    # match all M_1_hat and Gamma_hat corresponding Rf, matrix and scalar
    pattern = re.compile(
        r'Rf: (\d+).*?M_hat: \[\[(.*?)\], \[(.*?)\]\].*?Gamma_hat: ([\d\.Ee+-]+)',
        re.DOTALL
    )

    data = []
    for match in pattern.finditer(content):
        rf = int(match.group(1))
        row1 = list(map(float, match.group(2).split(',')))
        row2 = list(map(float, match.group(3).split(',')))
        M1 = np.array([row1, row2])
        gamma = float(match.group(4))
        data.append((rf, M1, gamma))

    # sort by Rf
    data.sort(key=lambda x: x[0])

    # store as numpy array list or stacked array
    rfs = np.array([d[0] for d in data])
    M1s = np.stack([d[1] for d in data], axis=0)  # shape: (n, 2, 2)
    gammas = np.array([d[2] for d in data])       # shape: (n,)

    return rfs, M1s, gammas


rfs, M1s, gammas = parse_file('bump_position_MLE_5.txt')
rfs = np.arange(1,25,2)
print("Rf:", rfs)
print("M1 shape:", M1s)
print("Gamma:", gammas)
# [-0.2518819272518158, -1.9999159574508667], [1.0002226829528809, -1.0002226829528809]
M1s_11 = M1s[:,0,0]
M1s_12 = M1s[:,0,1]
M1s_21 = M1s[:,1,0]
M1s_22 = M1s[:,1,1]


alpha_L = -(0.5*gammas**2)/(M1s_11+M1s_12)
print(alpha_L)
plt.figure()
plt.plot(rfs,alpha_L*(np.sqrt(np.array(2*np.pi))*0.5/40*rfs))
plt.xlabel('Rf')
plt.ylabel('alpha_L')
plt.savefig('alpha_L.png')
UE = []
US = []
for Rf in rfs:
    folder = "UeUs_outputs"
    filename = f"UeUs_Rf_{Rf:.2f}.npy"
    Ue = np.load(os.path.join(folder, filename))[:,0].mean(axis=0)
    Us = np.load(os.path.join(folder, filename))[:,1].mean(axis=0)
    UE.append(Ue)
    US.append(Us)
HE = []
for Rf in rfs:
    folder = "height_outputs"
    filename = f"height_Rf_{Rf:.2f}.npy"
    height = np.load(os.path.join(folder, filename))
    height = height.mean(axis=0)
    HE.append(height)
Lambda_list = []
for Rf in rfs:
    Lambda = (np.sqrt(np.array(2*np.pi))*0.5/40*Rf)
    Lambda_list.append(Lambda)
Lambda_list = np.array(Lambda_list)
HE = np.array(HE)
UE = np.array(UE)
US = np.array(US)
U_ES = M1s_12*UE
U_SE = M1s_21*US
U_EF = -(M1s_11+M1s_12)*UE
alpha_2 = M1s_11*HE
alpha_1 = -U_ES-(1-alpha_L)*U_EF
beta_E = -alpha_2*UE**-1*U_EF*(U_ES+(1-alpha_2)*U_EF)+US**-1*U_SE*(1-alpha_2)*U_EF
Lambda_p = (UE*beta_E)**-1*Lambda_list
# Plot Lambda_p
plt.figure(figsize=(3.5, 2.5))
plt.plot(rfs, Lambda_p)
plt.xlabel(r'$R_f$')
plt.ylabel(r'$\Lambda_p$')
plt.tight_layout()
plt.savefig('Lambda_p.eps', format='eps', bbox_inches='tight')
plt.savefig('Lambda_p.png', dpi=300, bbox_inches='tight')

inv_alpha2 = 1.0 / alpha_2         
alpha1_sq  = alpha_1 * alpha_1
T11 = M1s_11 - alpha_1 * inv_alpha2 * M1s_12
T12 = inv_alpha2 * M1s_12
T21 = alpha_1 * M1s_11 - alpha1_sq * inv_alpha2 * M1s_12 + alpha_2 * M1s_21 - alpha_1 * M1s_22
T22 = (alpha_1 * M1s_12 + alpha_2 * M1s_22) * inv_alpha2
# assemble
M_2 = np.stack([
    np.stack([T11, T12]),
    np.stack([T21, T22])
])
print(M_2.shape)
print(alpha_1.shape)
print(alpha_2.shape)
np.save('M_2_2.0.npy',M_2)
np.save('alpha_1_2.0.npy',alpha_1)
np.save('alpha_2_2.0.npy',alpha_2)
np.save('Lambda_p_2.0.npy',Lambda_p)
np.save('Sigma_E_2.0.npy',gammas)
with open('bump_position_MLE_5.txt', 'r') as f:
    f.write(f"Rf: {Rf}  M_2_hat: {M_2.tolist()}  "
                    f"alpha_1_hat: {alpha_1.tolist()}  "
                    f"alpha_2_hat: {alpha_2.tolist()}  "
                    f"Lambda_p_hat: {Lambda_p.tolist()}  "
                    f"Sigma_E_hat: {gammas.tolist()}\n")
f.close()
```
