## Supplementary for "Natural gradient Bayesian sampling automatically emerges in canonical cortical circuits": MLE_D_estimation.py

```
import torch
import torch.optim as optim
import numpy as np
import fit_lambda_p.HAM_cann_simulator as HAM
import os
import multiprocessing as mp

# ------------------------------------------------------------
# choose device once
# ------------------------------------------------------------
device = torch.device("cuda" if torch.cuda.is_available() else "cpu")
print("Using device:", device)

# ------------------------------------------------------------
# Step A:   direct estimator  U_SE
# ------------------------------------------------------------
def estimate_U_se(z_samples: torch.Tensor, dt: float):
    """z_samples : [T,2] or [B,T,2] on any device"""
    if z_samples.ndim == 3:
        z_S, z_E = z_samples[..., 1], z_samples[..., 0]
        num   = (z_S[:, 1:] - z_S[:, :-1]) / dt
        denom =  z_E[:, :-1] - z_S[:, :-1]
        U_se  = (num / (denom + 1e-12)).mean().item()
    else:
        z_S, z_E = z_samples[:, 1], z_samples[:, 0]
        U_se = (((z_S[1:] - z_S[:-1]) / dt) /
                (z_E[:-1] - z_S[:-1] + 1e-12)).mean().item()
    return U_se

# ------------------------------------------------------------
# Step B:   MLE for M11, M12, sigma_E
# ------------------------------------------------------------
def nll_zE(M11, M12, log_sigma, z_E, z_S, dt):
    mu   = z_E[..., :-1] + dt * (M11*z_E[..., :-1] + M12*z_S[..., :-1])
    diff = z_E[..., 1:] - mu
    var  = torch.exp(2*log_sigma) * dt
    sigma = torch.exp(log_sigma)
    return ((diff**2)/(2*var) + torch.log(sigma)).mean()

def fit_M11_M12_sigma(z_samples: torch.Tensor, dt: float,
                      lr=5e-2, epochs=1000, verbose=100):
    if z_samples.ndim == 3:        # [B,T,2]
        z_E, z_S = z_samples[..., 0], z_samples[..., 1]
    else:                          # [T,2]
        z_E = z_samples[:, 0][None, ...]
        z_S = z_samples[:, 1][None, ...]

    # parameters on the same device / dtype
    M11 = torch.tensor(0.1,  requires_grad=True, device=device)
    M12 = torch.tensor(-0.1, requires_grad=True, device=device)
    log_sigma = torch.tensor(-1.0, requires_grad=True, device=device)

    opt = optim.Adam([M11, M12, log_sigma], lr=lr)

    for ep in range(epochs):
        loss = nll_zE(M11, M12, log_sigma, z_E, z_S, dt)
        opt.zero_grad(); loss.backward(); opt.step()
        if verbose and ep % verbose == 0:
            print(f"ep {ep:4d}  NLL {loss.item():.5f}")

    sigma = torch.exp(log_sigma).item()
    return M11.item(), M12.item(), sigma

# ------------------------------------------------------------
# Convenience wrapper
# ------------------------------------------------------------
def estimate_drift_and_noise(z_samples: torch.Tensor, dt: float, verbose=True):
    z_samples = z_samples.to(device, dtype=torch.float32)

    U_se = estimate_U_se(z_samples, dt)
    if verbose: print(f"Estimated U_SE = {U_se:.4f}")

    M11, M12, sigma_E = fit_M11_M12_sigma(z_samples, dt, verbose=verbose)
    if verbose:
        print(f"M11={M11:.4f},  M12={M12:.4f},  sigma_E={sigma_E:.4f}")

    M_hat = torch.tensor([[M11,  M12],
                          [U_se, -U_se]], device="cpu")     # back to CPU
    Q_hat = torch.tensor([[sigma_E**2 * dt, 0.0],
                          [0.0,             0.0]], device="cpu")
    return M_hat, Q_hat, sigma_E, U_se

def one_bump_processing(Rf, seed):
    print(f"[Child PID {os.getpid()}] Start sim Rf={Rf}, seed={seed}")
    params = {
        'time_constant_exc': 1.0,
        'position_max': 180.0,
        'position_min': -180.0,
        'gaussian_width_exc': 40.0,
        'gaussian_width_ES': 20.0,
        'num_neurons': 180,
        'simulation_time': 10000.0,
        'time_step': 0.01,
        'recording_start': 20,
        'Fano_factor': 0.5,
        'normalization_k': 0.0005,
        'inhibitory_gain': 10,
        'input_position': 0,
        'feedforward_scale': 1.16429574032,
        't_steady': 20,
        'initial_mean_eq': 0,
        'initial_var_eq': 60,
        'initial_scale_eq': 1e-1
    }
    ZE, ZS, bump_height = HAM.bump_position_ham(params, Rf=Rf)
    ZE = ZE[2000:]
    ZS = ZS[2000:]
    print(ZE,ZS,np.var(ZE),np.var(ZS))

    print(f"[Child PID {os.getpid()}] Finished sim Rf={Rf}")
    return np.stack([ZE, ZS], axis=-1)  # shape: [T, 2]

# ------------------------------------------------------------
# main loop
# ------------------------------------------------------------

def main():
    N_TRIAL = 1000                          # generate 1000 trajectories for each R_f
    N_CPU   = mp.cpu_count()   
    dt = 0.01
    for Rf in range(25, 27, 2):
        fname = f"z_samples_Rf={Rf}.npy"
        if not os.path.exists(fname):
            print(f"File {fname} not found. Launching simulation for Rf={Rf}...")
            seeds = np.random.randint(0, 2**31-1, size=N_TRIAL)
            args  = [(Rf, s) for s in seeds]           # one_bump_processing(Rf,seed)

            with mp.Pool(processes=N_CPU) as pool:
                traj_list = pool.starmap(one_bump_processing, args)

            z_samples = np.stack(traj_list, axis=0)    # (N_TRIAL,T,2)
            np.save(fname, z_samples)
            print(f"[{Rf}]  saved trajectories to {fname}")
        else:
            result = np.load(fname)
            z_samples = torch.from_numpy(result).to(device)
        M_hat, Q_hat, sigma_E, U_se = estimate_drift_and_noise(z_samples, dt)
        print(f"Rf={Rf}, U_se={U_se:.4f}")

        with open("bump_position_MLE.txt", "a") as f:
            f.write(f"Rf: {Rf}  M_hat: {M_hat.tolist()}  "
                    f"Q_hat: {Q_hat.tolist()}  Gamma_hat: {sigma_E}\n")

if __name__ == "__main__":
    main()
```
