## Supplementary for "Natural gradient Bayesian sampling automatically emerges in canonical cortical circuits": readme.md

# $\Lambda_P$ Estimation

This repository provides the full pipeline for estimating the posterior kinetic precision $\Lambda_{P}$ at a range of feed-forward gain values $R_f$.

## Directory structure

```bash
.
├── height_outputs/# simulated trajectories and bump heights
├── long_trial_outputs/# simulated trajectories 
├── UeUs_outputs/# UeUs trajectories 
├── height.py        # compute bump-height H_E and H_S
├── UeUs.py          # compute U_E and U_S time-series
├── long_trial.py    # generate long reference trajectories
├── MLE_D_estimation.py  # stage-1 M-matrix + σ̂_E fit
├── lambdap.py       # analytic α₂, α₁, Λ_P evaluation + plot
└── run_all.sh           # convenience driver

```


## Quick start

A single command will reproduce every intermediate file and
 the final figure:

```bash
bash run_all.sh
```

The script executes all steps in the correct order and saves intermediate NumPy arrays plus the publication-quality plot
 `Lambda_p.png`. **We highly recommend you run code on a cluster!**

## Manual workflow

If you prefer to run the stages individually:

1. **Generate bump statistics**

   ```bash
   python scripts/height.py
   python scripts/UeUs.py
   python scripts/long_trial.py
   ```

   These scripts write `height_outputs/*.npy` and `UeUs_outputs/*.npy`.

2. **Fit the drift matrix $\widehat D$**

   ```bash
   python scripts/MLE_D_estimation.py
   ```

   Output is appended to `bump_position_MLE.txt`.

3. **Compute $\alpha_2,;\alpha_1,;\Lambda_{P}$ and plot**

   ```bash
   python scripts/lambdap.py
   ```

   The resulting plot is saved as `Lambda_p.png` (and `Lambda_p.eps` for LaTeX).

## Requirements

- Python ≥ 3.8
- PyTorch ≥ 1.11 (GPU optional)
- NumPy, SciPy, Matplotlib ≥ 3.5
