## Supplementary for "Natural gradient Bayesian sampling automatically emerges in canonical cortical circuits": run_all.sh

```
#!/usr/bin/env bash
#
# Reproduce the full Λ_P estimation pipeline
# ------------------------------------------
#   1.  height.py          → height_outputs/*.npy
#   2.  UeUs.py            → UeUs_outputs/*.npy
#   3.  long_trial.py      → z_samples_Rf=*.npy
#   4.  MLE_M1_estimation  → bump_position_MLE_5.txt
#   5.  lambdap.py         → Lambda_p.png / .eps
#
# Usage:
#   chmod +x run_all.sh
#   ./run_all.sh
#
# ------------------------------------------------------------------------------

set -euo pipefail

SECONDS=0
timestamp() { date +"%F %T"; }

echo "[$(timestamp)]  Λ_P pipeline started."

mkdir -p logs height_outputs UeUs_outputs

stage () {
  echo
  echo "[$(timestamp)]  === $* ==="
}

stage "1/5  Compute bump heights (H_E, H_S)"
python scripts/height.py      2>&1 | tee logs/height.log

stage "2/5  Compute U_E and U_S time-series"
python scripts/UeUs.py        2>&1 | tee logs/UeUs.log

stage "3/5  Generate long reference trajectories"
python scripts/long_trial.py  2>&1 | tee logs/long_trial.log

stage "4/5  Fit M₁ and σ_E via maximum likelihood"
python scripts/MLE_D_estimation.py \
                           2>&1 | tee logs/MLE_D.log

stage "5/5  Evaluate α₂, α₁, Λ_P and create figure"
python scripts/lambdap.py     2>&1 | tee logs/lambdap.log

echo
echo "[$(timestamp)]  Pipeline completed in $SECONDS seconds."
echo "Outputs:"
echo "  • Intermediate .npy files   → data/, height_outputs/, UeUs_outputs/"
echo "  • Drift/σ fit log           → bump_position_MLE_5.txt"
echo "  • Figure                    → Lambda_p.png  (+ .eps)"
echo "  • Verbose logs              → logs/*.log"
```
