## Supplementary for "Natural gradient Bayesian sampling automatically emerges in canonical cortical circuits": UeUs.py

```
import cann_simulator as HAM
import numpy as np
import multiprocessing as mp
import os
import re
import matplotlib.pyplot as plt
def Ue_Us_ham(args):
    params, Rf = args
    simulator = HAM.CANNSimulator(params)
    ZE, ZS, Ue, Us = HAM.bump_position_ham(params, Rf=Rf)
    return Rf,np.stack([Ue,Us],axis=-1)


if __name__ == "__main__":
    params = {
        'time_constant_exc': 1.0,
        'position_max': 180.0,
        'position_min': -180.0,
        'gaussian_width_exc': 40.0,
        'gaussian_width_ES': 20.0,
        'num_neurons': 180,
        'simulation_time': 1000.0,
        'time_step': 0.01,
        'recording_start': 20,
        'Fano_factor': 0.5,
        'normalization_k': 0.0005,
        'inhibitory_gain': 10,
        'input_position': 0,
        'feedforward_scale': 1.16429574032,
        't_steady': 20,
        'initial_mean_eq': 0,
        'initial_var_eq': 60,
        'initial_scale_eq': 1e-1
    }
    Rf_values = [r for r in range(1, 25,2)]
    args_list = [(params, Rf) for Rf in Rf_values]

    with mp.Pool(processes=10) as pool:
        results = pool.map(Ue_Us_ham, args_list)

    # save as dict
    Ue_Us_dict = {Rf: Ue_Us_array for Rf, Ue_Us_array in results}

    # save as numpy folder
    os.makedirs("UeUs_outputs", exist_ok=True)
    for Rf, arr in Ue_Us_dict.items():
        filename = f"UeUs_outputs/UeUs_Rf_{Rf:.2f}.npy"
        np.save(filename, arr)
        print(f"Saved {filename}, shape = {arr.shape}")
folder = "UeUs_outputs"
pattern = re.compile(r"UeUs_Rf_([0-9.]+)\.npy")

# read all files
Ue_Us_data = {}
for filename in os.listdir(folder):
    match = pattern.match(filename)
    if match:
        Rf = float(match.group(1))
        print(Rf)
        arr = np.load(os.path.join(folder, filename))
        print(arr)
        Ue_Us_data[Rf] = arr  # shape: (height_dim, time_len)

# sort
print(Ue_Us_data)
Rf_sorted = sorted(Ue_Us_data)
print(Rf_sorted)

plt.figure(figsize=(10, 6))
cmap = plt.cm.viridis
norm = plt.Normalize(vmin=min(Rf_sorted), vmax=max(Rf_sorted))
for Rf in Rf_sorted:
    Ue_Us_arr = Ue_Us_data[Rf]  # shape: (height_dim, time_len)
    U_e = Ue_Us_arr[:,0]
    # mean_height = height_arr.mean(axis=0)  # average across height_dim
    plt.plot(U_e, label=f"Rf={Rf:.2f}", alpha=0.5, c=cmap(norm(Rf)))
    plt.hlines(U_e.mean(), 0, len(U_e), colors=cmap(norm(Rf)))
plt.xlabel("Time step")
plt.ylabel("Ue")
plt.title("Ue over time for different Rf")
plt.legend()
plt.grid(False)
plt.tight_layout()
plt.savefig("Ue_over_time.png")

plt.figure(figsize=(10, 6))
cmap = plt.cm.viridis
norm = plt.Normalize(vmin=min(Rf_sorted), vmax=max(Rf_sorted))
for Rf in Rf_sorted:
    Ue_Us_arr = Ue_Us_data[Rf]  # shape: (height_dim, time_len)
    U_s = Ue_Us_arr[:,1]
    # mean_height = height_arr.mean(axis=0)  # average across height_dim
    plt.plot(U_s, label=f"Rf={Rf:.2f}", alpha=0.5, c=cmap(norm(Rf)))
    plt.hlines(U_s.mean(), 0, len(U_s), colors=cmap(norm(Rf)))
plt.xlabel("Time step")
plt.ylabel("Us")
plt.title("Us over time for different Rf")
plt.legend()
plt.grid(False)
plt.tight_layout()
plt.savefig("Us_over_time.png")
```
