## Supplementary for "Natural gradient Bayesian sampling automatically emerges in canonical cortical circuits": README.md

## Natural Gradient CANN Simulation

This repository provides Python scripts for simulating and analyzing Continuous Attractor Neural Networks (CANNs) in both one-dimensional (1D) and two-dimensional (2D) configurations. 

---

## Directory Structure

```bash
.
├── 1d/
│   ├── 1d.py                 # Main script for 1D CANN simulation
│   ├── cann_stimulator.py    # CANN class definition
│   ├── analysis_tool.py      # Analysis functions (KL divergence, cross-correlation, NG sampling)
│   └── plot_data.py          # Visualization tools for 1D data
├── 2d/
│   ├── 2d.py                 # Main script for 2D CANN simulation
│   ├── cannND.py             # cannND class definition (coupled 2D model)
│   ├── analysis_tool2D.py    # Analysis tools for 2D results
│   └── plot_data2D.py        # Plotting functions for 2D data
├── fit_lambda_p/
│   ├── height_outputs/# simulated trajectories and bump heights
│   ├── long_trial_outputs/# simulated trajectories 
│   ├── UeUs_outputs/# UeUs trajectories 
│   ├── height.py        # compute bump-height H_E and H_S
│   ├── UeUs.py          # compute U_E and U_S time-series
│   ├── long_trial.py    # generate long reference trajectories
│   ├── MLE_D_estimation.py  # stage-1 M-matrix + σ̂_E fit
│   ├── lambdap.py       # analytic α₂, α₁, Λ_P evaluation + plot
│   └── run_all.sh           # convenience driver
└── README.md                 # This file (example)
```

---

## Quick Start 🚀

For a streamlined execution, it is highly recommended to run the main simulation scripts for each dimension. These scripts are designed with sequential execution of specific code blocks in mind for optimal performance and time-saving.

**For 1D Simulation:**
Navigate to the `1d` directory and execute the lines within `1d.py` sequentially as indicated in the file.

**For 2D Simulation:**
Navigate to the `2d` directory and execute the lines within `2d.py` sequentially as indicated in the file.

**$\Lambda_P$ Estimation:**

A single command will reproduce every intermediate file and
 the final figure:

```bash
bash run_all.sh
```

The script executes all steps in the correct order and saves intermediate NumPy arrays plus the publication-quality plot
 `Lambda_p.png`. **We highly recommend you run code on a cluster!**.

**Note:** Detailed instructions for running specific parts of the simulations are commented within `1d.py` and `2d.py`.

---

## Manual Workflow for 1D & 2D CANN Simulation

If you prefer to understand or run specific components individually:

1.  **1D CANN Simulation & Analysis**
    * Navigate to the `1d/` directory.
    * **Run Simulation:** Execute the simulation parts of `1d.py`.
        * This utilizes `cann_stimulator.py` for the network model.
    * **Analyze Results:** Use functions from `analysis_tool.py` as called or adapted from `1d.py`.
    * **Visualize Data:** Employ `plot_data.py` for generating plots, typically invoked within `1d.py` or used separately with saved data.

2.  **2D CANN Simulation & Analysis**
    * Navigate to the `2d/` directory.
    * **Run Simulation:** Execute the simulation parts of `2d.py`.
        * This utilizes `cannND.py` for the 2D network model.
    * **Analyze Results:** Use functions from `analysis_tool2D.py` as called or adapted from `2d.py`.
    * **Visualize Data:** Employ `plot_data2D.py` for 2D visualizations, typically invoked within `2d.py` or used separately with saved data.


## Manual workflow for $\lambda_P$ estimation

If you prefer to run the stages individually:

1. **Generate bump statistics**

   ```bash
   python scripts/height.py
   python scripts/UeUs.py
   python scripts/long_trial.py
   ```

   These scripts write `height_outputs/*.npy` and `UeUs_outputs/*.npy`.

2. **Fit the drift matrix $\widehat D$**

   ```bash
   python scripts/MLE_D_estimation.py
   ```

   Output is appended to `bump_position_MLE.txt`.

3. **Compute $\alpha_2,;\alpha_1,;\Lambda_{P}$ and plot**

   ```bash
   python scripts/lambdap.py
   ```

   The resulting plot is saved as `Lambda_p.png` (and `Lambda_p.eps` for LaTeX).

## Requirements

- Python ≥ 3.8
- PyTorch ≥ 1.11 (GPU optional)
- NumPy, SciPy, Matplotlib ≥ 3.5
=======
##
